# Integrated single-cell (phospho-)protein and RNA detection uncovers phenotypic characteristics of human antibody secreting cells

**DOI:** 10.1101/2022.03.31.486501

**Authors:** Erik van Buijtenen, Wout Janssen, Paul Vink, Maurice J.M. Habraken, Laura J. A. Wingens, Andrea van Elsas, Wilhelm T.S. Huck, Jessie A.G.L. van Buggenum, Hans van Eenennaam

**Author notes:** These authors contributed equally to this work (Erik van Buijtenen, Wout Janssen). Shared senior author (Jessie A.G.L. van Buggenum, Hans van Eenennaam).

## Abstract

Antibody-secreting cells (ASCs) secrete IgM, IgA, or IgG antibodies and are key components of humoral immunity; however, little is known about unique characteristics of the Ig-classes due to limited availability of material and challenges to quantify many intracellular molecular modalities at a single-cell resolution. We combined a method to in vitro differentiate peripheral B-cells into ASCs with integrated multi-omic single-cell sequencing technologies to quantify subclass-specific hallmark surface markers, transcriptional profiles and signaling transduction pathway components. Our approach detected differential expression of plasmablast and plasma cell markers, homing receptors and IL-2, IL-6, JAK/STAT and mTOR signaling activity across Ig-subclasses. Taken together, our integrated multi-omics approach allowed high-resolution phenotypic characterization of single cells in a complex sample of in vitro differentiated human ASCs. Our strategy is expected to further our understanding of human ASCs in healthy and diseased samples and provide a valuable tool to identify novel biomarkers and potential drug targets.

**Teaser:** Integrated single-cell analysis allows tri-modal phenotypic analysis of in-vitro generated human antibody-secreting cells.

## Introduction

Upon activation, B-cells differentiate into plasma blasts and long-lived memory antibody-secreting cells (ASCs) (*1, 2*). An extraordinary number of control mechanisms, at both the cellular and molecular levels, underlie the generation and maintenance of plasmablasts and plasma cells from their B cell precursors (*3*). Aberrant plasma-blast and ASC activation/proliferation has been linked to several human diseases. In inflammatory and autoimmune diseases like systemic lupus erythematosus (SLE) or IgA nephropathy, aberrant ASC activation is linked to disease etiology and manifestation (*1*). In cancer, multiple myeloma (MM) is a malignancy derived from a plasma cell (PC) (*4*). A deeper insight into ASC phenotype can aid our understanding such diseases and identify biomarkers of disease and potential novel drug targets.

Human PCs only represent ∼0.25% of total bone marrow cells and home to different tissues in the human body, making extensive molecular analysis and functional studies challenging (*5*). Protocols for *in vitro* differentiation of human peripheral B-cells into ASCs (*6–8*) form the basis for phenotypic and functional studies (*9, 10*). Studies in mice have shown divergent gene expression profiles for the different Ig class PCs (*11*) and different expression profiles were observed comparing human class-switched germinal center (GC) B-cells from human tonsils as compared to non-class switched cells (*12*). However, bulk analysis obscures potential differences across ASCs secreting IgM, IgA, IgE or IgG antibodies. Single-cell technologies can characterize secreted antibody repertoire of individual cells (*13, 14*), and fully characterize the functional and transcriptional diversity of ASC (*15*).

Single-cell sequencing technologies have revolutionized our insight into the molecular phenotyping of many cell types of the human body. The combined quantification of mRNA and surface protein markers allows for additional (sub-)classification of cells (*16, 17*), while immune-detection by sequencing (ID-seq) (*18, 19*) and droplet-based QuRIE-seq (*20*) allows quantification of intracellular phospho-proteins. These methods rely on the reversible fixation of cells and immunostaining with DNA-tagged antibodies, enabling RNA sequencing and quantification of (phospho-)proteins in a single assay. Unfortunately, our fixation-based methods did not result in high-quality mRNA libraries in primary immune-cells (*20*). To circumvent this technical challenge, we combined multi-modal data from fixed and non-fixed cells. Using a common set of surface proteins we were able to integrate mRNA and intracellular (phospho-)proteins detection. Applying this method to *in vitro* differentiated human ASCs we identified distinctly different phenotypes of IgM, IgA and IgG expressing ASCs on both transcriptional and protein levels as well as active signal transduction.

## Results

### Phenotypic multi-modal single-cell analysis of human antibody secreting cells (ASCs)

We adopted an *in vitro* differentiation protocol of human peripheral B-cells to generate human ASCs (*6–8*). Isolated B-cells from healthy donors were stimulated with CpG for 1 day. Then, in three phases, activated B-cells were differentiated into ASCs using a mixture of cytokines and CD40L (Fig. 1a). Characterisation by flow cytometry showed that the procedure generated cells that displayed hallmarks of ASCs: IgD^-^CD27^++^, increased cell-size and gained expression of CD38 (*21*) (Fig. S1A-B). CD38+ ASCs showed increased mRNA expression levels for CD38, BLIMP1 and XBP1 as compared to naïve B-cells, as well as downregulation of proliferation marker Ki-67 and naïve B-cell specific transcription factor PAX-5 (Fig. S1 C). Consistent with ASC differentiation, Ab secretion was detected for all three Ig-classes from day 7 onwards and was most strongly increased on day 11 (Fig. S1D).

**Fig. 1.**
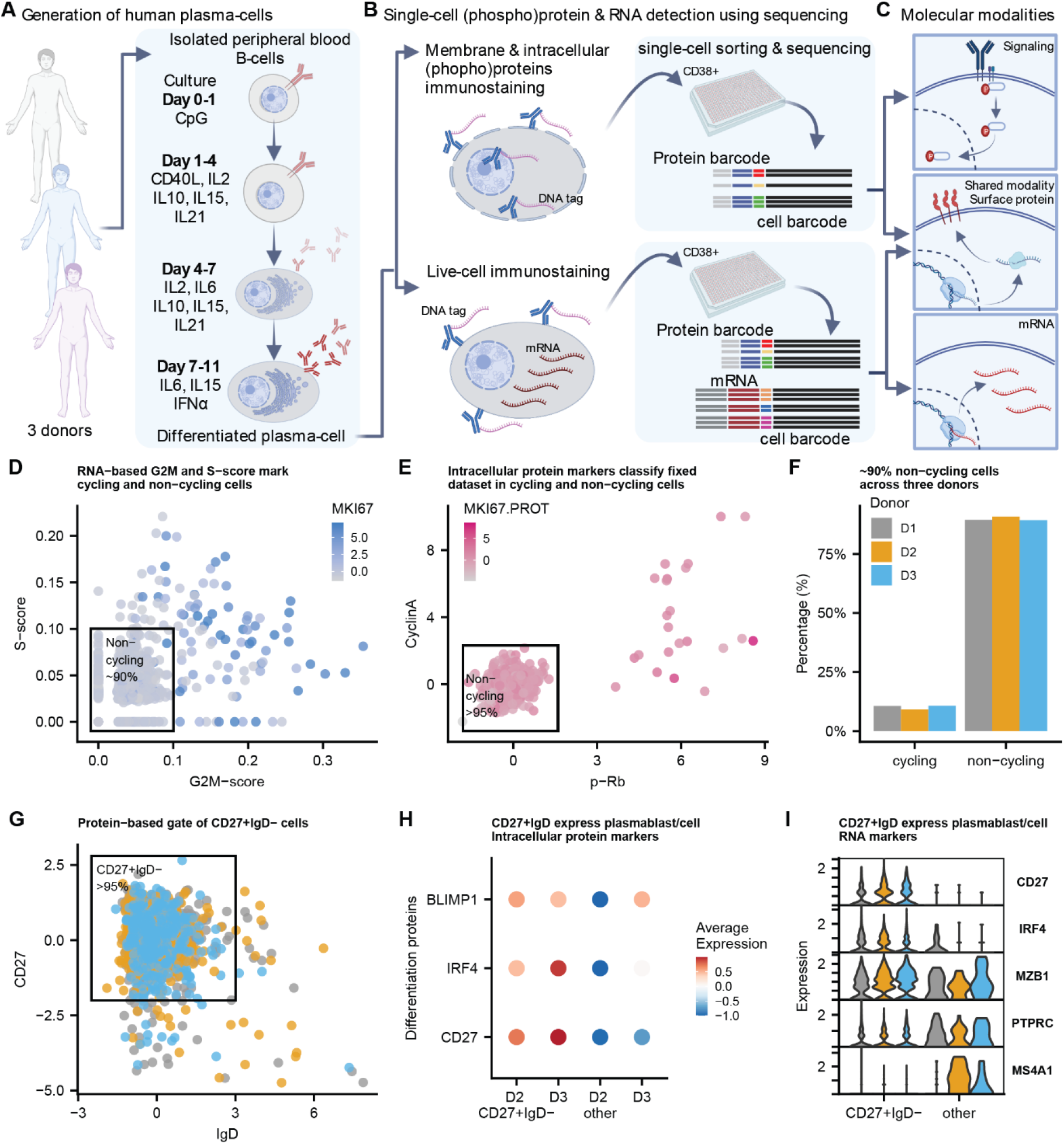
Experimental workflow for multi-modal single-cell analysis of in-vitro differentiated human antibody secreting cells (ASCs). (**A**) Peripheral B-cells isolated from healthy human blood are cultured in presence of indicated stimuli for 11 days. The protocol results in differentiated cells with ASC phenotype, capable of secreting antibodies. Created with BioRender.com (**B**) Day 11 cells are divided into one sample for fixation, permeabilization (allowing intracellular immune-staining with DNA-tagged antibodies) and a second sample for live-cell immunostaining with DNA-tagged antibodies. CD38+ single-cells are sorted into 384-well plates for multi-modal library preparation. (**C**) Analyses of both samples with a common set of surface-proteins, allows identification of molecular hallmarks: signaling (phospho-)proteins, protein levels and gene expression. Images (A-C) were created with BioRender.com. (**D**) RNA-based G2M-score and S-score based on pre-defined gene-sets is used for classification of each cells as non-cycling or cycling. Color represents normalized RNA-expression of proliferation marker *MKI67*. (**E**) Classification of non-cycling cells based on intracellular signal of p-Rb, Cyclin A and Ki67. (**F**) Percentage classified cycling or non-cycling cells per donor (N = 3). (**G**) CD27+/IgD-gate to select for differentiated PCs. Color represent three donors (legend see panel C). (**H**) Average (scaled) expression of selected intracellular proteins (**I**) Per-donor expression gene-expression of three differentiation markers (*CD27, IRF4* and *MZB1*) and two genes downregulated during PC differentiation (Wilcox, p-val < 0.05) in gated versus other cells (panel D). Colors represent donors (legend in panel C).

ASCs were collected on Day 11 and further purified by CD38+ cell-sorting (Fig. S2). These CD38+ ASCs were used for high resolution characterization via a single-cell multi-modal sequencing technology. To quantify signaling pathway activity (i.e. phosphorylation levels), surface markers, and mRNA expression on single-cell level, we developed a strategy based on CITE-seq (*16*), Immuno-Detection by Sequencing (ID-seq) (*18, 19*), RNA and Immuno-detection (RAID) (*22*) and droplet-based QuRIE-seq (*20*). To enable immunostaining and mRNA analysis on live ASCs for CITE-seq-like sample-preparation and immunostaining on fixed and permeabilized ASCs for intracellular immuno-detection by sequencing, we divided the ASCs from each donor into two portions before staining and sorting for multi-modal single-cell analysis (Fig 1B). Importantly, both samples were co-stained with a common set of 17 surface antibodies including IgM, IgA and IgG (Table S1-2). We reasoned that such a common ‘bridge modality’ should allow the integration and matching of the two parallel datasets based on the expression of these surface protein. Together, the workflow generates a dataset that includes information on ASC signaling pathway activity, protein levels and mRNA expression at single-cell resolution (Fig. 1C).

Quality control and filtering of the sequencing data (including > 300 genes per cell and < 5% mitochondrial gene counts per cell, sample specific miminum and maximum UMI counts per cell for the protein libraries, see Fig. S3 for full details) resulted in a selection of in total of 1,433 cells in the live-cell dataset and 1,038 cells in the fixed-cell dataset from three individual donors. To confirm the terminal differentiation of the *in-vitro* generated ASCs, we analyzed the cell-cycle state at both mRNA and protein-expression level. We categorized cells based on RNA-expression levels of cell-cycle genes, using the gene signature scoring method UCell (*23*) with G2M- and S-phase gene-lists as an input. Based on the computed scores and expression of proliferation marker *KI-67*, >95% of cells were classified as non-cycling (Fig. 1D), which was confirmed using intracellular protein expression of Cyclin A, p-Rb and Ki-67 (Fig. E). All three donors showed comparable percentages of dividing versus non-dividing cells (Fig. 1F, Fig. S4A).

ASCs, most specifically plasma blasts (PBs) and plasma cells (PCs), are characterized by IgD^-^CD27^+^CD38^+^ (*21*). Using the cell-sorted CD38+ cells, we compared IgD^-^CD27^+^ and IgD^+^CD27^-^ cells (Fig. 1G, Fig. S4B-C). CD27^+^IgD^-^ cells showed differential protein levels of ASC transcription factors BLIMP1 and IRF4 (Fig. 1H), which was further confirmed with the observed increased expression (Fig. S4 D-F) of *CD27, IRF4*, and *MZB1* genes (Fig. 1I) and downregulation of *PTPRC* (encoding CD45) and *MS4A1* genes (Fig. 1I) as compared to CD27^-^IgD^+^ cells. Taken together, our analyses confirmed that the in vitro procedure generated differentiated ASCs. For subsequent analysis, only non-proliferating IgD^-^CD27+CD38^+^ cells were studied.

### Single-cell mRNA and protein analysis reveals Ig-class specific phenotypes

We analysed the datasets using multi-omics factor analysis (MOFA+) (*24*). This analysis identifies shared variance across multiple molecular modalities, and allows integrated downstream analysis such as clustering. We supplied the model with four ‘modalities’: mRNA, fixed-protein, live-protein and common-protein, and grouped cells per donor to diminish donor-specific differences. (Fig. S5A). MOFA+ analysis computed 9 factors, of which factor 1 and 2 explain the most variance within the common modality (Supplementary Figure 5b). IgM, IgG and IgA proteins are the major contributing features to these factors, which suggests that factor 1 and 2 classify ASCs based on IgM, IgA and IgG levels (Figure 2A and Fig. S5 B-C). Protein expression of IgM, IgA and IgG (Fig 2A, upper panels in pink) correlated with the gene-expression levels of these Ig-types (Fig 2A, lower panels in blue). No variation of these factors was observed between the three individual donors or the live- and fixed-cell dataset (Fig. 2B, C, Fig. S5C). Based on this correlation, these factors were used for k-means clustering annotating individual cells as IgG, IgM or IgA subclass (Fig. S5D). Using an independent analysis method (PCA followed by clustering per modality) we confirmed the grouping of cells into Ig-subclasses (Fig. S6). Additionally we observe that all three donors displayed similar percentages of IgA, IgM and IgG ASCs (Fig. S7).

**Figure 2.**
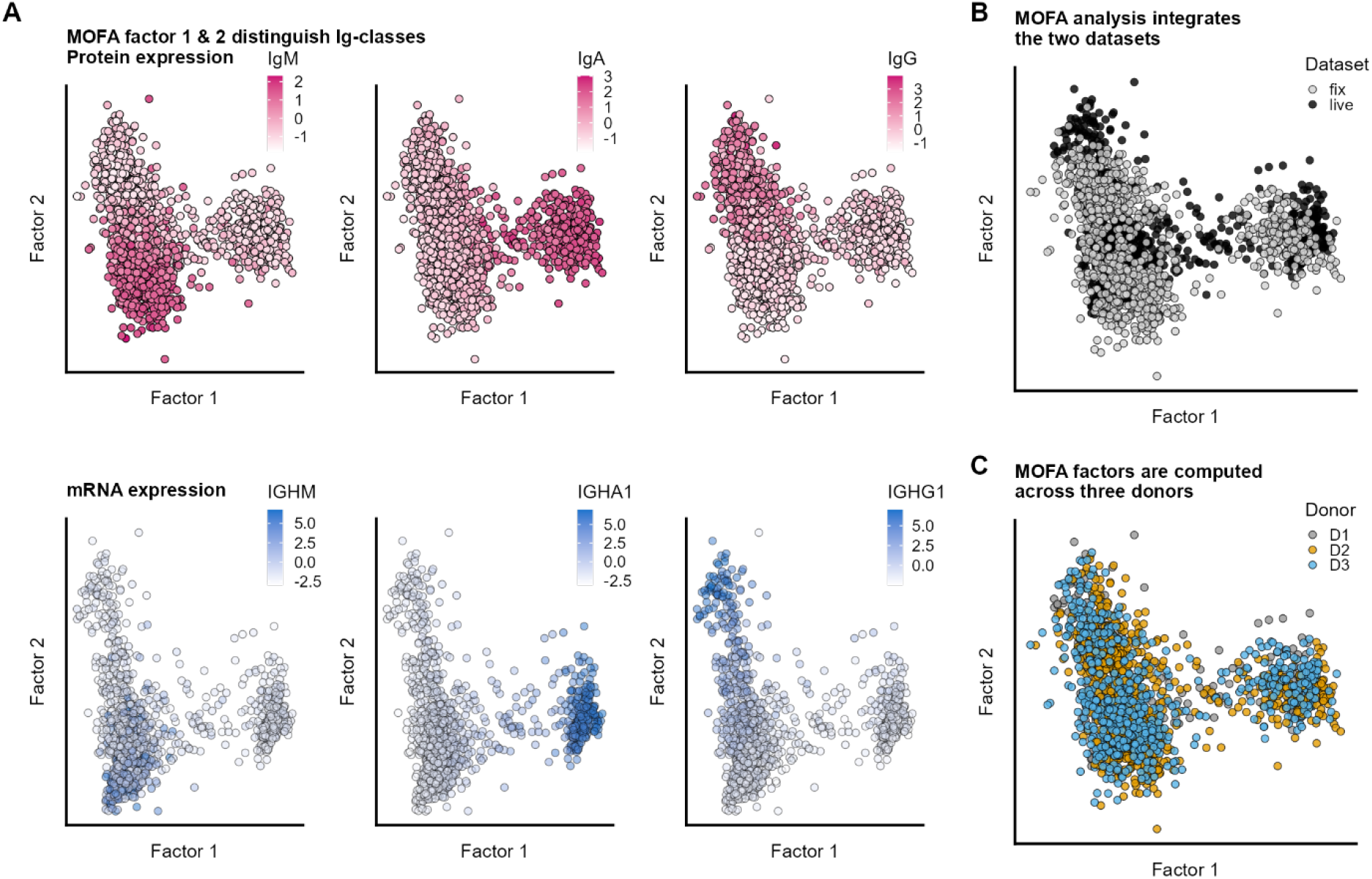
Joined analysis of all modalities using Multi-Omics Factor Analysis (MOFA+) reveals three mayor Ig-classes IgM, IgA and IgG across datasets and donors. (**A**) Protein-levels (upper panels, pink color) and gene-expression (lower panels, blue color) of IgM, IgA and IgG showing Factor 1 and Factor 2 split cells based on Ig-class. (**B**) MOFA factors 1 and 2 split sample independent of dataset origin (fix or live cell sample) (**C**) Factorplot coloured by donor shows no donor-effects.

**Figure 3.**
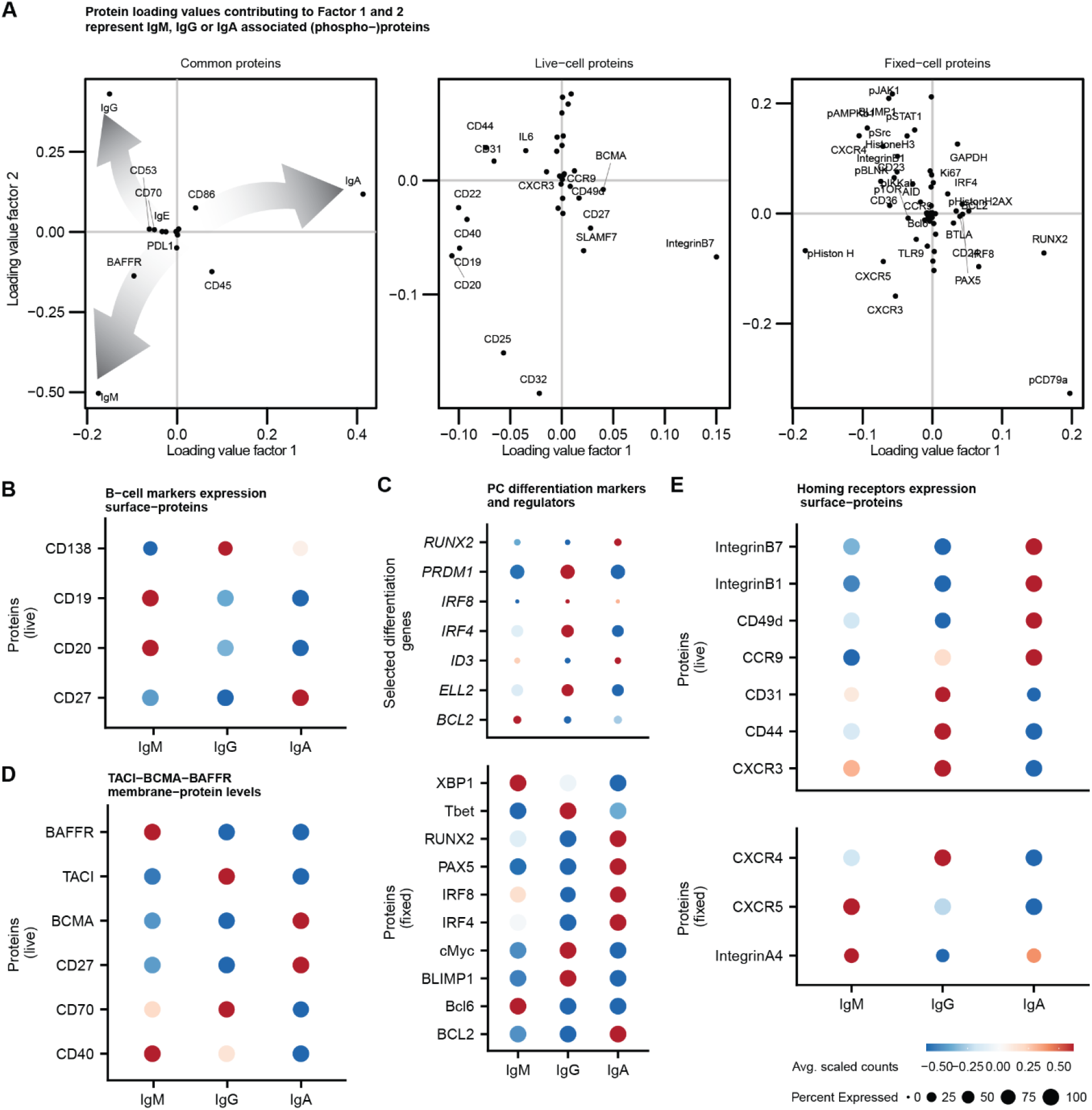
Exploration of protein and gene expression levels per Ig-classes IgM, IgA and IgG highlights subclass specific abundance of markers for B-cells, PC differentiation and homing-receptors. (**A**) Scatterplot of loading values per protein modality. Highlighted with text are selected contributing B-cell markers, differentiation related proteins and homing receptors. Grey arrows indicate direction of loadings contributing to the IgM, IgG or IgA subclass (**B**) Visualization of average expression of selected membrane proteins in IgM, IgG and IgA cells. (**C**) Manually curated list of intracellular differentiation genes and transcription factor proteins. (**D**) Scaled protein levels of BCMA, BAFFR and TACI. (**E**) Average expression of chemokine and adhesion related surface proteins. For all panels same legend applies presented as displayed in right lower corner.

**Figure 4.**
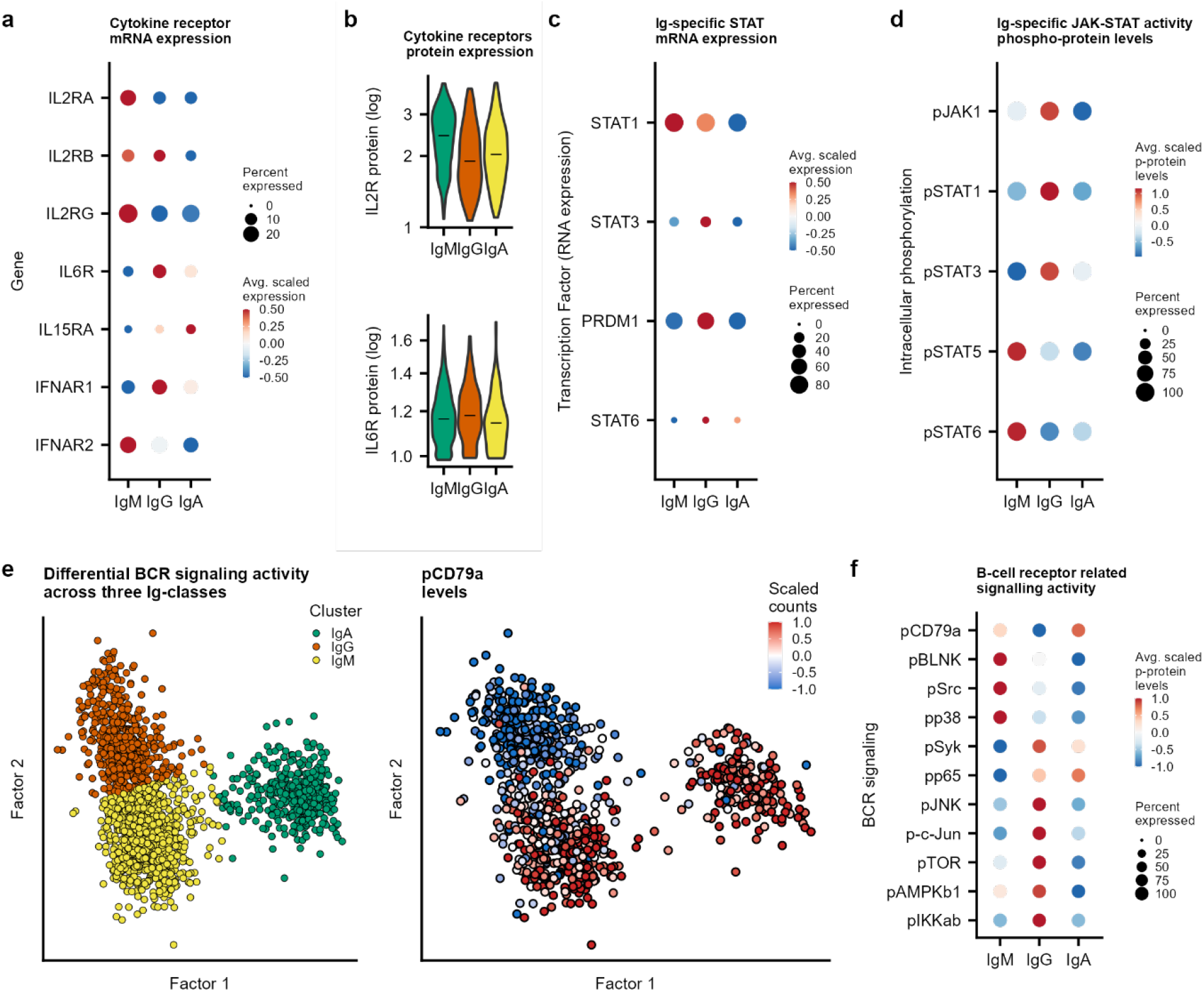
Ig-class specific cytokine, JAK-STAT, tonic B-cell receptor and mTOR signaling activity. (**A-B**) Gene (dotplot) and protein expression (violin plots) of cytokine receptors. (**C**) Gene expression of STAT-signaling components. (**D**) Average phosphorylation levels of JAK-stat signaling components. (**E**)UMAP representation of fixed dataset displaying annotated Ig-classes (left panel) and scaled phosphorylation levels of B-cell co-receptor CD79a (right panel). (**F**) Per sub-class average signaling activity of selected B-cell receptor, mTOR- and NFKb-related signaling components.

**Fig. 5.**
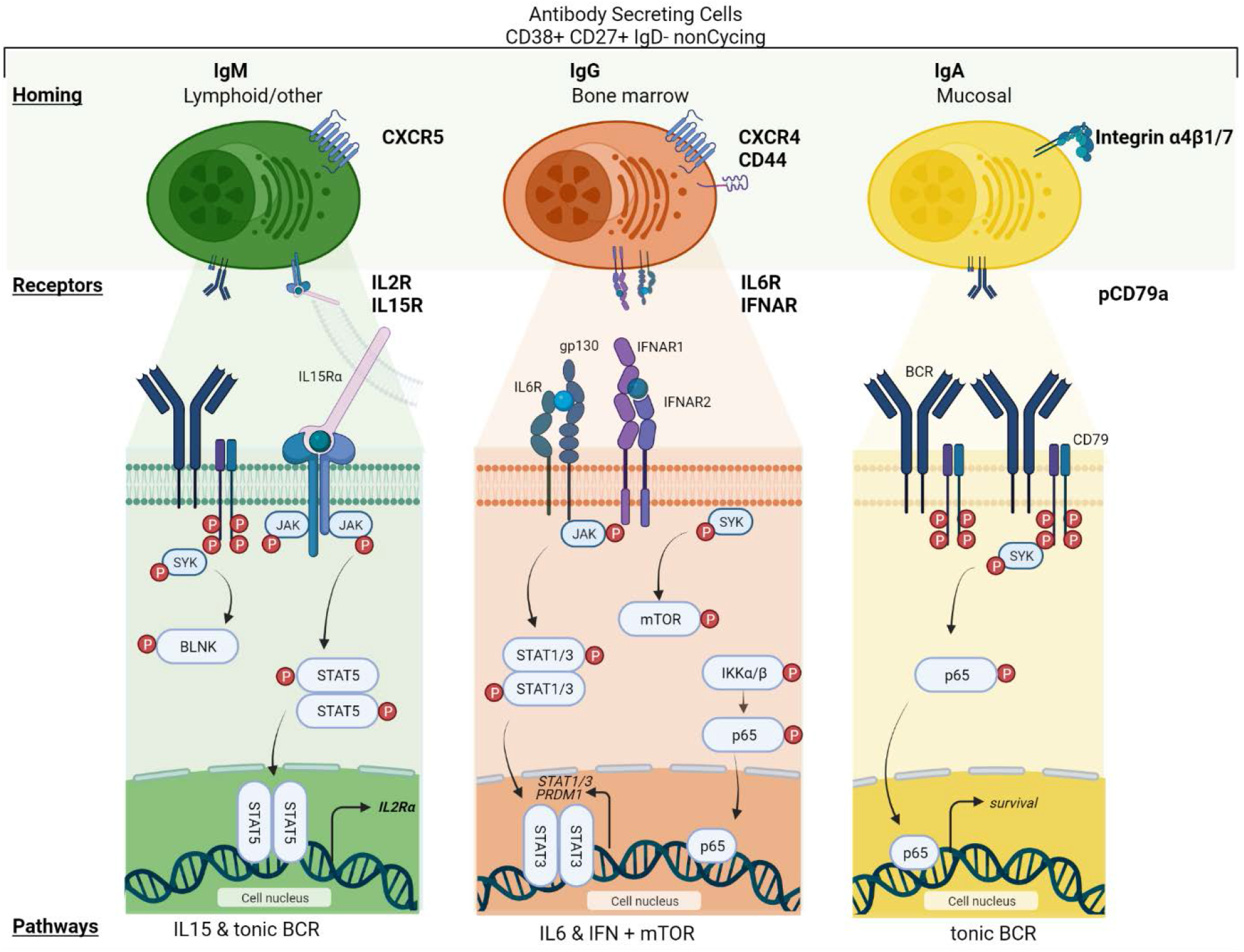
Highlights of human Ig-specific plasma-cells expression and signaling pathway activity. Single-cell multi-omics analysis revealed three major Ig-classes among the in-vitro differentiated human antibody secreting cells (ASCs): IgM (left), IgG (middle) and IgA (right). Each Ig-class shows a unique combination of phenotypes across multiple molecular modalities: surface-protein expression of homing and cytokine receptors, signal pathway activity and mRNA expression. Created with BioRender.com

To dissect phenotypic differences between ASCs secreting IgM, IgG and IgA, we identified proteins and genes with high loading values (weights) for factor 1 and 2, correlating with either IgA, IgG, or IgM (Figure 3A, Fig. S8). First, we studied the abundance of ASC markers across the different Ig-subclasses. All major population of human peripheral B cells can be identified using a small set of phenotypic markers: CD19, CD20, IgD, CD27, and CD38 (*21*). Using our analysis we only study IgD-CD27+ cells that were purified by CD38+ cell-sorting on Day 11 after in vitro differentiation of peripheral B-cells. As shown in Figure 3B, differences exist between the expression levels of these phenotypic markers across different Ig subclasses. Where CD19 and CD20 are found expressed most highly on IgM ASCs, CD19 and CD20 show low signals on IgG and IgA ASCs. Studying the expression of CD138, as a marker of more mature PC, it is interesting to note that only IgG ASCs highly express CD138, suggesting that IgG ASCs have a more PC phenotype, where IgM and IgA ASCs have a more PB phenotype. Also, when studying transcription and translation factors, IgG ASCs seem to reflect more strongly a PC phenotype with high levels of BLIMP1 on both mRNA (*PRDM1*) and protein level and *IRF4* on mRNA (Figure 3C). In accordance with literature, high levels of BLIMP1 coincide with low levels of PAX5 and Bcl6 protein. Interestingly, high levels of ell2 mRNA and c-myc and Tbet protein are also detected in the IgG ASCs as compared to IgA and IgM ASCs, where ell2 has been linked to IgG protein secretion in mice (*25*). In IgM ASCs, high levels of XBP1 protein are detected in absence of PAX5 (Figure 3C), where in IgA ASCs high levels of PAX5 repress XBP1 protein levels. The absence of BLIMP1 expression in both IgA and IgM ASCs and high BLC6 protein levels in IgM ASCs again suggests that IgM and IgA ASCs have a more PB phenotype. In IgA ASCs also other PAX5-regulated genes such as CD79A, IRF8 and ID3 were highly expressed (Fig. 3C) and high expression was observed for RUNX2, a crucial factor in IgA differentation of B-cells (*26*).

Different TNF-receptor family members have been implicated in B-cell and ASC differentiation. As shown in Figure 3D, highest expression of CD40 is found on IgM ASCs, consistent with its reported role in class-switching and survival (*27*). Expression of CD27 and its ligand CD70 appears to be mutual exclusive as described before, where high CD70 levels are observed on IgG ASCs as compared to IgM and IgA ASCs. Previously, CD70 signaling has been associated with specific downregulation of IgG, which is potentially explained by the differential expression of CD70 on IgG ASCs (*28*). Comparing expression of the BCMA, TACI and BAFF-R indicates that BCMA is most highly expressed on IgA ASCs, while TACI is most highly expressed on IgG ASC and BAFF-R is highest expressed on IgM ASCs (Figure 3D). These observations are very well aligned with the observed defects in Ig production seen in TACI-/-knockout mice (*29*) and low IgM and IgG levels in the BAFF-R-/-knockouts (*30*).

IgG PCs are thought to migrate to the bone marrow, while IgA PCs migrate to mucosal associated lymphoid tissue (*2*). We next studied whether different Ig-subclasses of ASCs are differently imprinted to mediate migration to these sites. Indeed, the bone marrow homing receptor CXCR4 was highly expressed on IgG ASCs of which the ligand, CXCL12 is expressed by bone marrow stromal cells (Fig. 3E) (*31*). Furthermore, CD44 and CD31 were highest expressed on IgG ASCs, which have been shown to be important for adhesion to the bone marrow stroma (Fig. 3E) (*32*). In contrast to IgG ASCs, IgA ASCs expressed integrins α4 (CD49d) and β1 and β7, which have been reported to direct IgA PCs to respiratory and intestine mucosal tissue, respectively (Fig. 3E) (*33*). IgM PCs can remain in the lymphoid organs or migrate to sites of infection or mucosal tissue, the latter supported by the moderate expression of integrins α4 and β7 (Fig. 3E) and CXCR5 (under direction of CD40) on IgM ASCs (*34*). Overall, our single-cell method revealed that ASCs expressing different isotype-classes, differentially express transcriptional factors, membrane proteins and homing receptors.

### Ig-subclasss show differential activity of cytokine signaling across modalities

To establish the sensitivity for cytokine-induced signaling across IgM, IgA and IgG ASCs, we analyzed receptor expression, intracellular phospho-proteins and downstream gene expression of IL2/IL15, IL6, JAK/STAT and IFNA pathways. As shown in Figure 4, IgM ASCs show elevated expression of *IL-2RA, IL-2RB, IL-2RG* mRNA and high-affinity IL2 receptor (CD25) protein (Fig. 4A-B) as well as moderate levels of pJAK1 and high levels of pSTAT5/6 (Fig. 4C-D). Signal transduction is most likely driven by IL-15 as from Day 7 no IL-2 was added to the *in vitro* culture, and IL-15 is present between Day 7-11 (Fig. 4A-D) (*35*). In contrast, IgG ASCs express high levels of *IL-6R* mRNA as well as the corresponding protein IL-6R (Fig. 4A). IL-6 signaling induces high levels of p-STAT3, which in literature is described as a key inducer of *PRDM1* expression (*36*). In addition, high expression of IFNAR1 and IFNAR2 protein was detected in IgG ASCs (Fig. 4AB) and IFN-α signaling mediators were most abundantly detected in IgG ASCs with pJAK1, pSTAT1 and *STAT1* gene expression (Fig. 4C-D). Finally, IgA ASCs showed low JAK/STAT pathway activation and modest IL-6R and STAT3/6 expression, suggesting IgA PCs are less dependent or sensitive to these cytokines. Taken together, our analysis allowed us to observe two distinctly different cytokine-induced signaling profiles in PCs: IgM shows high IL-15 and STAT5/6 signaling, while IgG shows high IL-6/IFNR and STAT1/3 signaling (Figure 4D).

### Tonic BCR signaling and mTOR show different activity in IgM, IgG or IgA cells

Differentiation of B cells in ASCs/PCs is accompanied by a switch from membrane to secreted Ig, thereby losing expression of the BCR and its signal transduction. Recent studies in both mice and human PCs revealed that IgM and IgA PCs still express a BCR capable of signaling, suggesting these cells can still detect and respond to antigens (*37, 38*). To study whether evidence of active BCR signaling can be detected in the *in vitro* differentiated ASCs, we analyzed different elements of BCR pathway activation. Both IgA and IgM ASCs had high levels of phosphorylated CD79a, the immediate downstream signal transduction protein of the BCR (Fig. 4F). Also other downstream signal components of the BCR, pSRC, pBLNK and pp38, were detected in IgM ASCs, while pSYK was detected in IgA ASCs (Fig. 4F). In contrast to IgM and IgA ASCs, and in accordance with recent reports, little or no activation of BCR pathway was dectected in IgG ASCs (Fig. 4F). In fact, we observed phosphorylation of Syk, mTOR and observed NF-κB pathway activation as detected by pIKK and pP65, suggestive of an active BCR-independent signal transduction in IgG ASCs. Reversely, no or limited activation of NF-κB and/or mTOR is detected in IgA or IgM ASCs.

## Discussion

In this study, we employed an *in vitro* differentiation protocol to obtain large quantities of human ASCs from peripheral B-cells. Our multimodal single cell analysis method detected in this heterogenous sample of in vitro differentiated non-cycling IgD^-^ CD27^++^CD38^+^ ASCs the Ig-subclass specific expression of B cells phenotypic markers, transcription factors, TNF-receptor mediated survival receptors and expression of niche-specific homing receptors. More importantly, our method allowed the detection of active cytokine-induced signal transduction and B-cell receptor (BCR) and mTOR/AKT signaling that demonstrated that each Ig-subclass has differential sensitivity towards these stimuli based on the differential expression of their receptors.

Antibody secreting cells, composed of plasma-blasts and plasma-cells are crucially important to maintain circulating immunoglobulin levels and provide long-lasting immunity by their sometimes life-long survival in dedicated niches in the body. Aberrant PCs have been linked to human diseases like rheumatoid arthritis, SLE, IgA nephropathy and multiple myeloma. A detailed molecular phenotyping and deep understanding of human ASC differentiation and PC survival is important to understand the etiology of such diseases as well as to identify molecular targets for drug discovery.

Recently, studies using single-cell RNA sequencing technologies have provided more understanding on the (heterogeneity in) phenotype of aberrant PCs (*39–42*). However, these studies only provide insight in the mRNA expression of different cells, whereas in our study, we combine mRNA, extracellular and intracellular (phospho-)protein information at the single cell level. One of the current major challenges in the single-cell field is integrating datasets across samples and molecular modalities. Computational strategies encounter several considerations as how to define anchors, scalability and handling missing data (*43*). Several of these challenges are being addressed by recently developed tools including MOFA+ (*24*), multiVI (*44*), COBOLT (*45*), StabMap (*46*) scMVP (*47*), and Bridge Integration (*48*). So-far there was no illustration of integrating mRNA datasets with a comprehensive intracellular phospho-protein and transcription factor dataset using a common set of surface proteins. Here, we illustrate the succesfull use of the MOFA+ framework bridging single-cell mRNA quantification with detailed insight in signal pathway activity from the same cell (sub)type. In principle, as long as the phenotype (such as cell subtypes in this manuscript) are being quantified in the shared molecular modality, any other associated phenotypes in separate datasets can be integrated and explored. Taken together, based on our analyses performed, integration of both datasets allowed us to gain detailed insight into human ASCs across three molecular modalities.

Both in human and mice, IgM and IgA but not IgG ASCs have been shown to express a functional BCR, which can activate intracellular signaling (*37, 38*), and tonic BCR signaling has been reported for pre- and naive B-cells as an important survival signal (*49*). The in vitro differentiated IgM and IgA ASCs show all hallmarks of tonic BCR signaling, suggesting that these cells still rely on this pathway for survival. It has been proposed that CD45, a positive regulator of BCR signaling, triggers receptor tyrosine kinases to phosphorylate CD79a and CD79b, thereby initiating tonic signaling. This could explain the tonic signaling observed in this culture, as CD45 expression, was higher in IgM and IgA cells as compared to IgG cells (*49*). Also the observed expression of BCL6 in IgM ASCs may positively affect tonic BCR signaling in B-cell lymphoma cells (*50*). In contrast, IgG ASCs displayed no p-CD79a or downstream signaling, but our methods did reveal strong phosphorylation of SYK and mTOR pathway components. Apart from BCR induced activation, SYK can be activated through different ITAM containing receptors (*51*) and SYK may regulate activation of the mTOR pathway supporting contributing to cell growth and proliferation (*52, 53*). Additionally, in IgG we detected active IL-6 signaling and a NF-κB signature. This signaling axis, in concert with the BLIMP1-XBP1 axis, has been described to induce mTOR activity, and antibody production through regulating the unfolded protein response (*20, 54*). Taken together with the phenotypic markers (i.e. CD138, PAX5, ell2 etc.) using our multi-modal omics approach, we were able, in a single biological experiment, to reveal that the obtained IgG ASCs are defined by active IL-6, JAK-STAT, SYK and mTOR signaling and carry all hallmarks of IgG PCs.

The rapidly expanding range of single-cell technologies provides quantitative information about different ‘layers’ in the cell, but often not on multiple layers at the same time. We have demonstrated that integrating single-cell datasets on mRNA, surface markers and intracellular (phoshpo-)protein via the quantification of a shared modality is a powerful strategy to characterize the transcriptional wiring and signaling pathway activity in in-vitro differentiated primary cells. Our strategy yielded valuable insight into the molecular differences across modalities between human IgM, IgA and IgG ASCs. We envision that an integrated single-cell multi-omic analysis pipeline is especially suitable for studying cells in the PBMCs compartment, where the cell-type specific response to a range of stimuli provides a deeper understanding of key processes in immunology or blood borne cancers.

## Materials and Methods

### Antibody labeling

Carrier-free antibodies (Table S1-2) were conjugated to 5’-Azide-oligos (Table S3-4, Biolegio, The Netherlands) as described in Supplementary text. In brief, dibenzocyclooctyne-S-S-N-hydroxysuccinimidyl-ester functionalized antibodies were incubated with 3x molar access azide-oligo for 16 hours at 4°C. Unconjugated oligos were removed using 100K amicon centrifuge filters (Merck, USA).

### B-cell isolation and culture

Total B-cells were isolated from Buffy coat from healthy volunteers with written consent and ethics comity approval (Sanquin, the Netherlands) as described in Supplementary text. Subsequent differentiation cultures were performed in Iscove’s Modified Dulbecco’s Medium (Thermo Fisher Scientific, USA) supplemented with 10% FBS (Thermo Fisher Scientific, USA) and 1% Penicillin-Streptomycin (Thermo Fisher Scientific, USA). Purified B-cells were differentiated for 11 days as visualized in Figure 1A and described in detail in Supplementary text.

### Antibody staining and cell sorting

Day 11 cells were harvested and split in two tubes, one to be fixed and permeabilized and one for live cells analysis. Cells were immediately fixed by adding 2x concentrated fixative to the cell suspension (5 mM dithiobis (succinimidyl propionate) (DSP, Thermo Fisher Scientific, USA) and 5 mM succinimidyl 3-(2-pyridyldithio)propionate (SPDP, Thermo Fisher Scientific, USA) in PBS (Thermo Fisher Scientific, USA). Cells were incubated at RT for 45 minutes while gently shaking. Fixed cells were quenched and permeabilized with 100 mM Tris-HCl pH 7.5, 150 mM NaCl and 0.1% Triton X100 (Thermo Fisher Scientific, USA) for 10 minutes at RT. Cells were washed once and blocked for 45 minutes in 0.5X protein-free blocking buffer (Thermo Fisher Scientific, USA) with 0.2 mg/ml dextran sulfate (Sigma Aldrich, USA) and 0.5 U/ml RNasin plus (Promega, USA) in PBS (Thermo Fisher Scientific, USA). Cells were stained in blocking buffer containing DNA-tagged antibodies (Table S3) and CD38-PE-Cy7 (Table S5) for one hour at RT. Following staining, cells were washed twice with the blocking buffer and re-suspended in PBS containing 5 mg/ml BSA (Thermo Fisher Scientific, USA), 0.5 U/ml RNsing plus (Promega, USA) and 0.1μg/ml DAPI (Biolegend, USA).

Non-fixed live cells were were immediately washed twice in ice cold blocking buffer. Cells were stained in ice cold blocking buffer containing DNA-tagged Abs (Table S4) and CD38-PE-Cy7 Ab (Table S5). Cells were incubated on ice in the dark for 20 min. Cells were washed three times in ice cold blocking buffer and re-suspended in blocking buffer containing 0.1 μg/ml DAPI (Biolegend, USA).

Using the FACS Melody (BD Biosciences, USA), cells were sorted single-cell in 384-wells PCR plates (Biorad, USA) containing 100 nl water containing 7.5 ng/μl unique Cellseq-2 primers (Table S6) and 5 μl miniral oil (Sigma Aldrich, USA). Plates were stored at -80 °C until further use. Oligonucleotide sequences were adapted to allow sequencing of the transcripts/ARC sequences in read 1 and the cell barcode and UMI in read 2 (Table S4).

### Library preparation

Plates with sorted cells were thawed on ice. For fixed samples, cells were reverse-crosslinked, releasing RNA and antibody-barcodes as described by Gerlach et.al (*22*). Live-cells were mearly incubated for 5 minutes at 65 degrees. Reverse transcription was performed as described by Gerlach et.al (*22*). After pooling cells, the mRNA was separated from antibody barcodes using 0.6x AMPure XP machnetic beads (Beckman Coulter, USA). Then, mRNA sequencing libraries were generated as described by Gerlach et.al. (*22*) using primers listed in Table S4. From the separated antibody barcodes sample, protein libraries were generated as described in detail in Supplementary text. mRNA and protein libraries were sequenced with NextSeq500.

### Data analysis

Bcl2fasq software (illumina) was used for data-demultiplexing. The CEL-seq2 pipeline (*55*) and CITE-seq-count (*56*) tool were used to create count tables of mRNA or protein libraries respectively. Standard settings were used with some minor modifications. The CEL-seq2 pipeline was run using settings [min_bc_quality = 10, cut_length = 50]. Additionally, the read names for read 1 and read 2 were swapped to allow compatibility of our barcode design with the pipeline. The CITE-seq-count settings were kept at default with exeption of [ max_error of Ab barcode = 1, umi_collpasing_dist = 1].

Extensive documentation of quality control, filtering (Supplementary Figure 3 and data-normalization and processing can be found at https://vanbuggenum.github.io/Multimodal-Plasmacell_manuscript/QC.html). In brief, cells were filtered based on either mRNA library quality (Livecell dataset: > 300 genes, < 5% mitochondrial counts per cell) or protein library (>1500 and <9000 protein counts, > 40 different proteins per cell); Fixed dataset: >2500 and <20000 protein counts, > 65 proteins per cell). RNA-counts were normalized using ‘single-cell transform’ (Seurat V4) (*57*). Then, normalized counts were scaled, while regressing out differences in % mitochondrial counts and total-counts. Protein counts were normalized using the CLR method. (*57*) Then, the normalized protein-counts were scaled with regressing out variance in number of proteins detected and total-counts.

### Multi-modal analysis

To identify non-cycling ASCs, first a cell-cycle scoring for S-phase and G2M-phase was calculated using Ucell algorithm (*23*) using gene lists provided by the Seurat R-package (V4) (*57*). For the live dataset, a gate was determined based on these two scores. For the fixed dataset, p-Rb and CyclinA were used to identify cycling cells instead of mRNA-based cell-cycle score. To determine differentiated phenotype, protein levels of CD27 and IgD were used for gating. These non-cycling CD27+IgD-cells were kept for further analysis using MOFA+ (*24*). In brief, cells were grouped per donor, and the model received four molecular modalities. PCA analysis was performed using normalized and scaled counts of Ig-proteins.

## Acknowledgments

We thank Klaas Mulder for providing 384-well plates with Celseq2 oligo’s, Marijke Baltissen for assistance with sequencing, and Dyah Karjosukarso for processing FASTQ files generating the count tables.

## Funding

This study was supported (in part) by research funding from Aduro Biotech (EvB, WJ, PV, MC, AE and HvE)

Radboud University (LJAW, WTSH and JAGLvB)

Spinoza Grant (WTSH)

VENI grant from the Netherlands Organisation for Scientific Research VI.Veni.202.228 (JAGLvB)

## Author contributions

EvB, WJ, PV, and MJMH established and performed the in-vitro differentiation protocol. EvB and WJ performed the flow cytometry, qPCR and ELISA experiments. EvB and LJAW performed the multi-modal sequencing experiment. JAGLvB conceived and performed the data-integration and analyzed the sequencing results. AE and HE conceived the project, WTSH nd HE supervised the project, EvB, JAGLvB, WTSH and HvE wrote the manuscript. All authors approved the final version of the manuscript.

## Data and materials availability

Detailed code documentation of the analysis and generation of figures is available from https://vanbuggenum.github.io/Multimodal-Plasmacell_manuscript/. The fastq files and count are accessible through GEO (54) Series accession number GSE189953 (https://www.ncbi.nlm.nih.gov/geo/query/acc.cgi?acc=GSE189953).

## References

1. F. Hiepe, A. Radbruch, Plasma cells as an innovative target in autoimmune disease with renal manifestations. Nat. Rev. Nephrol. 12, 232–240 (2016).

2. J. Tellier, S. L. Nutt, Plasma cells: The programming of an antibody-secreting machine. Eur. J. Immunol. 49, 30–37 (2019).

3. S. L. Nutt, P. D. Hodgkin, D. M. Tarlinton, L. M. Corcoran, The generation of antibody-secreting plasma cells. Nat. Rev. Immunol. 15, 160–171 (2015).

4. N. W. C. J. van de Donk, C. Pawlyn, K. L. Yong, Multiple myeloma. Lancet. 397, 410–427 (2021).

5. L. W. M. M. Terstappen, S. Johnsen, I. M. J. Segers-Nolten, M. R. Loken, Identification and characterization of plasma cells in normal human bone marrow by high-resolution flow cytometry. Blood. 76, 1739–1747 (1990).

6. M. Jourdan, H. De Boussac, E. Viziteu, A. Kassambara, J. Moreaux, In vitro differentiation model of human normal memory B cells to long-lived plasma cells. J. Vis. Exp. 2019, 58929 (2019).

7. M. Jourdan, A. Caraux, J. De Vos, G. Fiol, M. Larroque, C. Cognot, C. Bret, C. Duperray, D. Hose, B. Klein, An in vitro model of differentiation of memory B cells into plasmablasts and plasma cells including detailed phenotypic and molecular characterization. Blood. 114, 5173–5181 (2009).

8. R. Itoua Maïga, G. Bonnaure, J. Tremblay Rochette, S. Néron, Human CD38hiCD138+ plasma cells can be generated in vitro from CD40-activated switched-memory B lymphocytes. J. Immunol. Res. 2014 (2014), doi:10.1155/2014/635108.

9. M. Jourdan, M. Cren, N. Robert, K. Bolloré, T. Fest, C. Duperray, F. Guilloton, D. Hose, K. Tarte, B. Klein, IL-6 supports the generation of human long-lived plasma cells in combination with either APRIL or stromal cell-soluble factors. Leukemia. 28, 1647–1656 (2014).

10. A. Kassambara, L. Herviou, S. Ovejero, M. Jourdan, C. Thibaut, V. Vikova, P. Pasero, O. Elemento, J. Moreaux, RNA-sequencing data-driven dissection of human plasma cell differentiation reveals new potential transcription regulators. Leukemia. 35, 1451–1462 (2021).

11. M. J. Price, S. L. Hicks, J. E. Bradley, T. D. Randall, J. M. Boss, C. D. Scharer, IgM, IgG, and IgA Influenza-Specific Plasma Cells Express Divergent Transcriptomes. J. Immunol. 203, 2121–2129 (2019).

12. H. W. King, N. Orban, J. C. Riches, A. J. Clear, G. Warnes, S. A. Teichmann, L. K. James, Single-cell analysis of human B cell maturation predicts how antibody class switching shapes selection dynamics. Sci. Immunol. 6 (2021), doi:10.1126/sciimmunol.abe6291.

13. K. Eyer, R. C. L. Doineau, C. E. Castrillon, L. Briseño-Roa, V. Menrath, G. Mottet, P. England, A. Godina, E. Brient-Litzler, C. Nizak, A. Jensen, A. D. Griffiths, J. Bibette, P. Bruhns, J. Baudry, Single-cell deep phenotyping of IgG-secreting cells for high-resolution immune monitoring. Nat. Biotechnol. 35, 977–982 (2017).

14. A. Gérard, A. Woolfe, G. Mottet, M. Reichen, C. Castrillon, V. Menrath, S. Ellouze, A. Poitou, R. Doineau, L. Briseno-Roa, P. Canales-Herrerias, P. Mary, G. Rose, C. Ortega, M. Delincé, S. Essono, B. Jia, B. Iannascoli, O. Richard-Le Goff, R. Kumar, S. N. Stewart, Y. Pousse, B. Shen, K. Grosselin, B. Saudemont, A. Sautel-Caillé, A. Godina, S. McNamara, K. Eyer, G. A. Millot, J. Baudry, P. England, C. Nizak, A. Jensen, A. D. Griffiths, P. Bruhns, C. Brenan, High-throughput single-cell activity-based screening and sequencing of antibodies using droplet microfluidics. Nat. Biotechnol. (2020), doi:10.1038/s41587-020-0466-7.

15. M. Broketa, P. Bruhns, Single-Cell Technologies for the Study of Antibody-Secreting Cells. Front. Immunol. 12, 1–9 (2022).

16. M. Stoeckius, C. Hafemeister, W. Stephenson, B. Houck-Loomis, P. K. Chattopadhyay, H. Swerdlow, R. Satija, P. Smibert, Simultaneous epitope and transcriptome measurement in single cells. Nat. Methods. 14, 865–868 (2017).

17. V. M. Peterson, K. X. Zhang, N. Kumar, J. Wong, L. Li, D. C. Wilson, R. Moore, T. K. McClanahan, S. Sadekova, J. A. Klappenbach, Multiplexed quantification of proteins and transcripts in single cells. Nat. Biotechnol. 35, 936–939 (2017).

18. J. A. G. van Buggenum, J. P. Gerlach, S. E. J. Tanis, M. Hogeweg, P. W. T. C. Jansen, J. Middelwijk, R. van der Steen, M. Vermeulen, H. G. Stunnenberg, C. A. Albers, K. W. Mulder, Immuno-detection by sequencing enables large-scale high-dimensional phenotyping in cells. Nat. Commun. 9, 2384 (2018).

19. R. A. P. M. van Eijl, J. A. G. L. van Buggenum, S. E. J. Tanis, J. Hendriks, K. W. Mulder, Single-Cell ID-seq Reveals Dynamic BMP Pathway Activation Upstream of the MAF/MAFB-Program in Epidermal Differentiation. iScience. 9, 412–422 (2018).

20. F. Rivello, E. van Buijtenen, K. Matuła, J. A. G. L. van Buggenum, P. Vink, H. van Eenennaam, K. W. Mulder, W. T. S. Huck, Single-cell intracellular epitope and transcript detection reveals signal transduction dynamics. Cell Reports Methods. 1, 100070 (2021).

21. J. L. Halliley, C. M. Tipton, J. Liesveld, A. F. Rosenberg, J. Darce, I. V. Gregoretti, L. Popova, D. Kaminiski, C. F. Fucile, I. Albizua, S. Kyu, K.-Y. Chiang, K. T. Bradley, R. Burack, M. Slifka, E. Hammarlund, H. Wu, L. Zhao, E. E. Walsh, A. R. Falsey, T. D. Randall, W. C. Cheung, I. Sanz, F. E.-H. Lee, Long-Lived Plasma Cells Are Contained within the CD19-CD38hiCD138+ Subset in Human Bone Marrow. Immunity. 43, 132–145 (2015).

22. J. P. Gerlach, J. A. G. van Buggenum, S. E. J. Tanis, M. Hogeweg, B. M. H. Heuts, M. J. Muraro, L. Elze, F. Rivello, A. Rakszewska, A. van Oudenaarden, W. T. S. Huck, H. G. Stunnenberg, K. W. Mulder, Combined quantification of intracellular (phospho-)proteins and transcriptomics from fixed single cells. Sci. Rep. 9, 1469 (2019).

23. M. Andreatta, S. J. Carmona, bioRxiv, in press, doi:10.1101/2021.04.13.439670.

24. R. Argelaguet, D. Arnol, D. Bredikhin, Y. Deloro, B. Velten, J. C. Marioni, O. Stegle, MOFA+: a statistical framework for comprehensive integration of multi-modal single-cell data. Genome Biol. 21, 111 (2020).

25. K. S. Park, I. Bayles, A. Szlachta-McGinn, J. Paul, J. Boiko, P. Santos, J. Liu, Z. Wang, L. Borghesi, C. Milcarek, Transcription Elongation Factor ELL2 Drives Ig Secretory-Specific mRNA Production and the Unfolded Protein Response. J. Immunol. 193, 4663–4674 (2014).

26. K. Watanabe, M. Sugai, Y. Nambu, M. Osato, T. Hayashi, M. Kawaguchi, T. Komori, Y. Ito, A. Shimizu, Requirement for Runx Proteins in IgA Class Switching Acting Downstream of TGF-$β$1 and Retinoic Acid Signaling. J. Immunol. 184, 2785–2792 (2010).

27. T. Kawabe, T. Naka, K. Yoshida, T. Tanaka, H. Fujiwara, S. Suematsu, N. Yoshida, T. Kishimoto, H. Kikutani, The immune responses in CD40-deficient mice: Impaired immunoglobulin class switching and germinal center formation. Immunity. 1, 167–178 (1994).

28. R. Arens, M. A. Nolte, K. Tesselaar, B. Heemskerk, K. A. Reedquist, R. A. W. van Lier, M. H. J. van Oers, Signaling through CD70 Regulates B Cell Activation and IgG Production. J. Immunol. 173, 3901–3908 (2004).

29. G. T. Mantchev, C. S. Cortesão, M. Rebrovich, M. Cascalho, R. J. Bram, TACI Is Required for Efficient Plasma Cell Differentiation in Response to T-Independent Type 2 Antigens. J. Immunol. 179, 2282–2288 (2007).

30. C. R. Smulski, H. Eibel, BAFF and BAFF-receptor in B cell selection and survival. Front. Immunol. 9, 1–10 (2018).

31. A. Radbruch, G. Muehlinghaus, E. O. Luger, A. Inamine, K. G. C. Smith, T. Dörner, F. Hiepe, Competence and competition: The challenge of becoming a long-lived plasma cell. Nat. Rev. Immunol. 6, 741–750 (2006).

32. G. Cassese, S. Arce, A. E. Hauser, K. Lehnert, B. Moewes, M. Mostarac, G. Muehlinghaus, M. Szyska, A. Radbruch, R. A. Manz, Plasma Cell Survival Is Mediated by Synergistic Effects of Cytokines and Adhesion-Dependent Signals. J. Immunol. 171, 1684–1690 (2003).

33. E. J. Kunkel, E. C. Butcher, Plasma-cell homing. Nat. Rev. Immunol. 3 (2003), pp. 822–829.

34. C. Wei, Y. Chen, L. Xu, B. Yu, D. Lu, Y. Yu, Z. Lei, R. Tang, S. Zhou, J. Zhu, X. Chen, C. Su, CD40 Signaling Promotes CXCR5 Expression in B Cells via Noncanonical NF-κ B Pathway Activation. J. Immunol. Res. 2020, 1–6 (2020).

35. A. M. Ring, J. X. Lin, D. Feng, S. Mitra, M. Rickert, G. R. Bowman, V. S. Pande, P. Li, I. Moraga, R. Spolski, E. Özkan, W. J. Leonard, K. Christopher Garcia, Mechanistic and structural insight into the functional dichotomy between IL-2 and IL-15. Nat. Immunol. 13, 1187–1195 (2012).

36. S. A. Diehl, H. Schmidlin, M. Nagasawa, S. D. van Haren, M. J. Kwakkenbos, E. Yasuda, T. Beaumont, F. A. Scheeren, H. Spits, STAT3-Mediated Up-Regulation of BLIMP1 Is Coordinated with BCL6 Down-Regulation to Control Human Plasma Cell Differentiation. J. Immunol. 180, 4805 (2008).

37. D. Pinto, E. Montani, M. Bolli, G. Garavaglia, F. Sallusto, A. Lanzavecchia, D. Jarrossay, A functional BCR in human IgA and IgM plasma cells. Blood. 121, 4110–4114 (2013).

38. P. Blanc, L. Moro-Sibilot, L. Barthly, F. Jagot, S. This, S. De Bernard, L. Buffat, S. Dussurgey, R. Colisson, E. Hobeika, T. Fest, M. Taillardet, O. Thaunat, A. Sicard, P. Mondière, L. Genestier, S. L. Nutt, T. Defrance, Mature IgM-expressing plasma cells sense antigen and develop competence for cytokine production upon antigenic challenge. Nat. Commun. 7, 1–14 (2016).

39. A. Gulla, K. C. Anderson, Multiple myeloma: The (r)evolution of current therapy and a glance into the future. Haematologica. 105, 2358–2367 (2020).

40. J. S. Jang, Y. Li, A. K. Mitra, L. Bi, A. Abyzov, A. J. van Wijnen, L. B. Baughn, B. Van Ness, V. Rajkumar, S. Kumar, J. Jen, Molecular signatures of multiple myeloma progression through single cell RNA-Seq. Blood Cancer J. 9 (2019), doi:10.1038/s41408-018-0160-x.

41. Y. C. Cohen, M. Zada, S. Y. Wang, C. Bornstein, E. David, A. Moshe, B. Li, S. Shlomi-Loubaton, M. E. Gatt, C. Gur, N. Lavi, C. Ganzel, E. Luttwak, E. Chubar, O. Rouvio, I. Vaxman, O. Pasvolsky, M. Ballan, T. Tadmor, A. Nemets, O. Jarchowcky-Dolberg, O. Shvetz, M. Laiba, O. Shpilberg, N. Dally, I. Avivi, A. Weiner, I. Amit, Identification of resistance pathways and therapeutic targets in relapsed multiple myeloma patients through single-cell sequencing. Nat. Med. 27, 491–503 (2021).

42. H. Zeng, L. Wang, J. Li, S. Luo, Q. Han, F. Su, J. Wei, X. Wei, J. Wu, B. Li, J. Huang, P. Tang, C. Cao, Y. Zhou, Q. Yang, Single-cell RNA-sequencing reveals distinct immune cell subsets and signaling pathways in IgA nephropathy. Cell Biosci. 11, 203 (2021).

43. R. Argelaguet, A. S. E. Cuomo, O. Stegle, J. C. Marioni, Computational principles and challenges in single-cell data integration. Nat. Biotechnol. 39, 1202–1215 (2021).

44. T. Ashuach, M. I. Gabitto, M. I. Jordan, N. Yosef, MultiVI: deep generative model for the integration of multi-modal data. bioRxiv (2021), doi:10.1101/2021.08.20.457057.

45. B. Gong, Y. Zhou, E. Purdom, Cobolt: integrative analysis of multimodal single-cell sequencing data. Genome Biol. 22, 351 (2021).

46. S. Ghazanfar, C. Guibentif, J. C. Marioni, StabMap: Mosaic single cell data integration using non-overlapping features. bioRxiv (2022), doi:10.1101/2022.02.24.481823.

47. G. Li, S. Fu, S. Wang, C. Zhu, B. Duan, C. Tang, X. Chen, G. Chuai, P. Wang, Q. Liu, A deep generative model for multi-view profiling of single-cell RNA-seq and ATAC-seq data. Genome Biol. 23, 20 (2022).

48. Y. Hao, T. Stuart, M. Kowalski, S. Choudhary, P. Hoffman, A. Hartman, A. Srivastava, G. Molla, S. Madad, C. Fernandez-granda, R. Satija, Dictionary learning for integrative, multimodal, and scalable single-cell analysis (2022).

49. J. G. Monroe, ITAM-mediated tonic signalling through pre-BCR and BCR complexes. Nat. Rev. Immunol. 6 (2006), pp. 283–294.

50. P. Juszczynski, L. Chen, E. O’Donnell, J. M. Polo, S. M. Ranuncolo, R. Dalla-Favera, A. Melnick, M. A. Shipp, BCL6 modulates tonic BCR signaling in diffuse large B-cell lymphomas by repressing the SYK phosphatase, PTPROt. Blood. 114, 5315–5321 (2009).

51. A. Mócsai, J. Ruland, V. L. J. Tybulewicz, The SYK tyrosine kinase: a crucial player in diverse biological functions. Nat. Rev. Immunol. 2010 106. 10, 387–402 (2010).

52. S. Fruchon, S. Kheirallah, T. Al Saati, L. Ysebaert, C. Laurent, L. Leseux, J. J. Fournié, G. Laurent, C. Bezombes, Involvement of the Syk-mTOR pathway in follicular lymphoma cell invasion and angiogenesis. Leukemia. 26, 795–805 (2012).

53. L. Leseux, S. M. Hamdi, T. Al Saati, F. Capilla, C. Recher, G. Laurent, C. Bezombes, Syk-dependent mTOR activation in follicular lymphoma cells. Blood. 108, 4156–4162 (2006).

54. S. Benhamron, S. P. Pattanayak, M. Berger, B. Tirosh, mTOR Activation Promotes Plasma Cell Differentiation and Bypasses XBP-1 for Immunoglobulin Secretion. Mol. Cell. Biol. 35, 153–166 (2015).

55. T. Hashimshony, N. Senderovich, G. Avital, A. Klochendler, Y. de Leeuw, L. Anavy, D. Gennert, S. Li, K. J. Livak, O. Rozenblatt-Rosen, Y. Dor, A. Regev, I. Yanai, CEL-Seq2: sensitive highly-multiplexed single-cell RNA-Seq. Genome Biol. 17, 77 (2016).

56. P. R. and bbimber and B. F. and santiagorevale and G. Gui, Hoohm/CITE-seq-Count: 1.4.2. Zenodo (2019), doi:10.5281/zenodo.2590196.

57. Y. Hao, S. Hao, E. Andersen-Nissen, W. M. Mauck, S. Zheng, A. Butler, M. J. Lee, A. J. Wilk, C. Darby, M. Zager, P. Hoffman, M. Stoeckius, E. Papalexi, E. P. Mimitou, J. Jain, A. Srivastava, T. Stuart, L. M. Fleming, B. Yeung, A. J. Rogers, J. M. McElrath, C. A. Blish, R. Gottardo, P. Smibert, R. Satija, Integrated analysis of multimodal single-cell data. Cell, 1–15 (2021).

