## Supplementary Material for "Integrated single-cell (phospho-)protein and RNA detection uncovers phenotypic characteristics of human antibody secreting cells"

**This PDF file includes:**

Supplementary Text
Figs. S1 to S8
Tables S1 to S5

**Supplementary Text**

Detailed description of antibody functionalization and conjugation

Antibodies were buffer exchanged into 0.2M NaHCO3 (pH 8.3) (Sigma Aldrich, USA) using Zeba™ Spin Desalting Columns, 40K MWCO (Thermo Fisher Scientific, USA) and aftefunctionalized with dibenzocyclooctyne-S-S-N-hydroxysuccinimidyl ester (Sigma Aldrich, USA) (10x molar axcess, 2h at room temperature (RT)). Functionalized antibodies were buffer exchanged into PBS (Thermo Fisher Scientific, USA) using 30K amicon centrifuge units (Merck, USA). Subsequently, 5' Azide modified oligos (Table S3-4, Biolegio, The Netherlands) were added to 25 or 50 µg antibody in 3x molar excess and incubated for 16 hours at 4°C. Ab-DNA conjugets were buffer exchanged into PBS (Thermo Fisher Scientific, USA) containing 0.05 % sodium azide (Sigma Aldrich, USA) and 0.1 mM EDTA (Sigma Aldrich, USA), removing exces oligonucleotides, using 100K amicon centrifuge filters (Merck, USA). Labelling efficiency was determined by non-reducing SDS-PAGE gel electrophoresis.

Detailed description of B-cell isolation and differentiation culture

B-cells were pre-enriched using the RossetteSep Human B-cell Enrichment cocktail according manufacturer's instructions (StemCell, USA). After Ficoll density gradient total B-cells were further isolated using the B cell isolation Kit II according manufacturer's instructions (Miltenyi Biotec, Germany). Purity of isolated B cells was determined by flow cytometry and was >99% CD19+ and CD20+.

5*10E6 purified B-cells were seeded in 500 µl medium in 14 ml round bottom tubes (VWR, the Netherlands) and stimulated for 16 hours 37°C/5 % CO2 with CpG ODN2006, Class B (1 µg/ml, Invivogen, USA) by adding 500 µl medium containing 2 µg/ml CpG.
Cells were washed once by adding 4 ml of medium and centrifuging cells at 200 g for 5 min and removing supernatant completely. Then cells were incubated for 3 days at 37°C/5 % CO2 in 5 ml fresh medium containing 500 ng/ml rhCD40L (Miltenyi Biotec, Germany), 20 U/ml rhIL-2 (Sigma Aldrich, USA), 50 ng/ml rhIL-10 (R&D Systems, USA), 10 ng/ml rhIL-15 (R&D Systems, USA), 100 ng/ml rhIL-21 (Miltenyi Biotec, Germany). Then, cells were grown another 3 days at 37°C/5 %CO2 with 5 ml of new medium containing 20 U/ml rhIL-2, 50 ng/ml rhIL-10, 10 ng/ml rhIL-15, 100 ng/ml rhIL-21 and 50 ng/ml rhIL-6 (R&D Systems, USA). Finally, cells were incubated 4 days at 37°C/5 % CO2 in 5 ml fresh medium containing 10 ng/ml rhIL-15, 50 ng/ml rhIL-6 and 500 U/ml rhINFα (R&D Systems, USA).

Detailed library preparation protocol of antibody-barcodes

After size separation of the protein and mRNA libraries, the protein library was purified with 1.5x AMPure XP magnetic beads and eluted in 10 µL of nuclease-free water. Subsequently, the sample was PCR amplified after prior determination of the number of cycles needed by qPCR: 0.5 µL of purified material was mixed with 6.5 µL nuclease-free water, 10 µL of 2x Kapa HiFi HotStart PCR mix (Roche, Switzerland), 1 µL of 20X EvaGreen Dye (Thermo Fisher Scientific, USA), and 2 µL of REV unique indexing primer and the Nextera primer (Table S4) primer mix (5 µM each) (Biolegio, The Netherlands). Then the PCR reaction was started by adding: 0.5 µL of nuclease-free water, 12.5 µL of 2X Kapa HiFi HotStart PCR mix (Roche, Switzerland), and 2.5 µL of REV unique indexing primer and the Nextera primer (Table S4) mix (5 µM each)

(Biolegio, The Netherlands) to the 9.5 μL of the sample. The PCR thermal cycling was as follows: 2 minutes at 95 °C, followed by x cycles of (x = number previously determined by qPCR): 30 seconds at 95 °C, 30 seconds at 64 °C and 40 seconds at 72 °C, and terminated with 5 minute-incubation at 72 °C. PCR amplified product was purified by addition of 1.2X AMPure XP magnetic beads, eluted in 10 μl water and its quality was verified by ds-DNA concentration measurement with Qubit ds-DNA High Sensitivity assay (Thermo Fisher Scientific, USA) and Bioanalyzer (Agilent 2100, USA) characterization. If the quality of the samples was high enough, the protein libraries were sequenced together with the mRNA libraries on with NextSeq500 (targeting 50 million reads per sample).

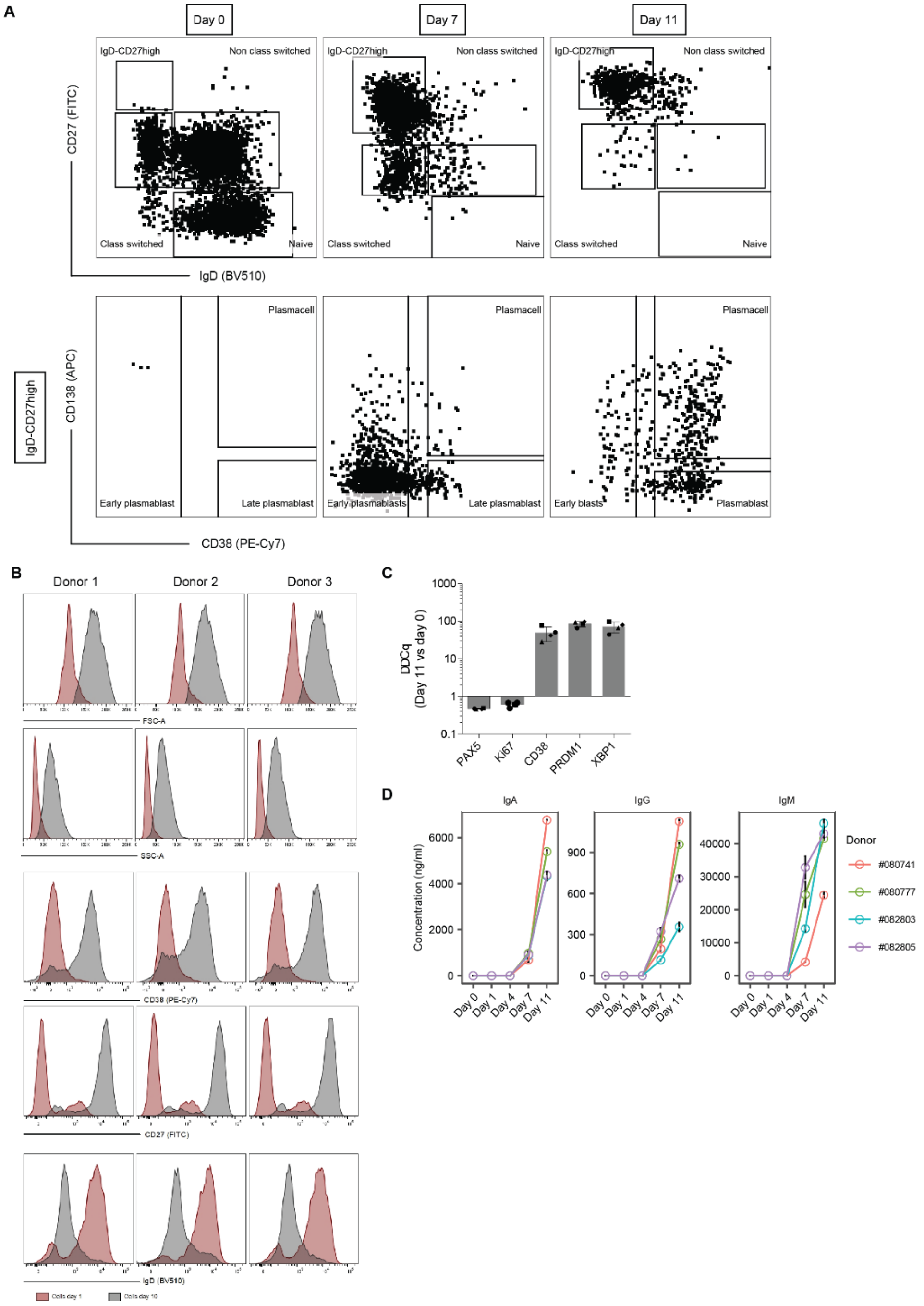


**Fig. S1. Molecular validation of the *in-vitro* differentiation of B-cells into plasma-cells.**

(**A**) Cytometry analysis of CD27 and IgD at 0, 7 or 11 days after differentiation. Indicated gates highlight types of cells during differentiation of B-cells. (**B**) Flow cytometry analysis of cells on day 0 and day 10 with analysis of cell size (FSC-A, SSC-A), of proteins up-regulated during differentiation CD38 and CD27 or expression profiles of protein downregulated during differentiation: IgD. (**C**) qPCR analysis of cells sorted at day 11 compared to day 0: *CD38*, *BLIMP1* and *XBP1*, *Ki67* and *PAX-5*. (**D**) ELISA results of IgM, IgA and IgG detected in culture supernatant after day 7. Ab secretion increased at day 11 of the culture, confirming the generation of Ab secreting cells.

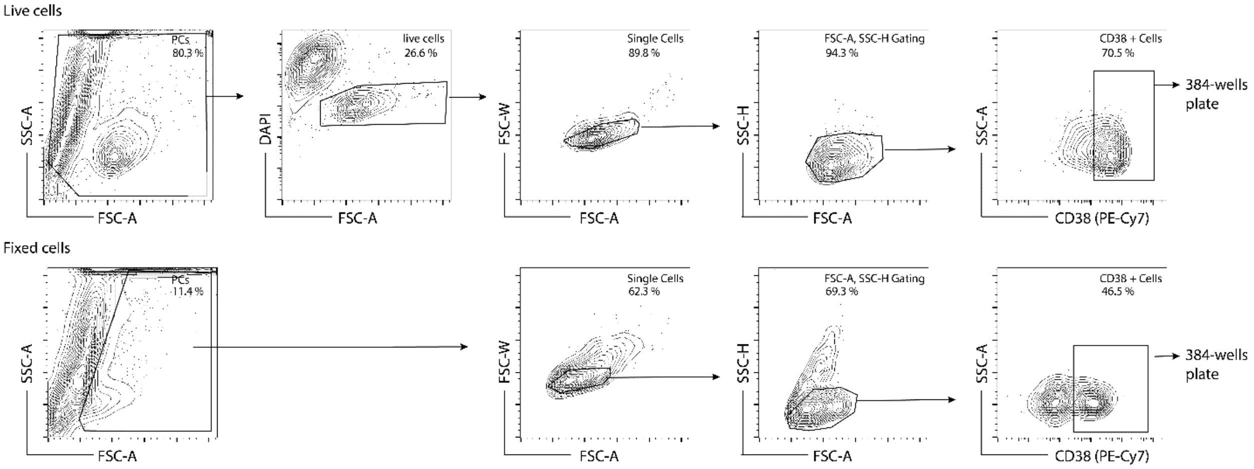


**Fig. S2. Sorting strategy of live (upper panels) or fixed (lower panels) cells.**

Live cells were gated on DAPI- to exclude dead cells. CD38+ cells were sorted in 384 wells plates. Fixed cells were gated on single-cells and CD38+ cells were sorted in 384 well plates.

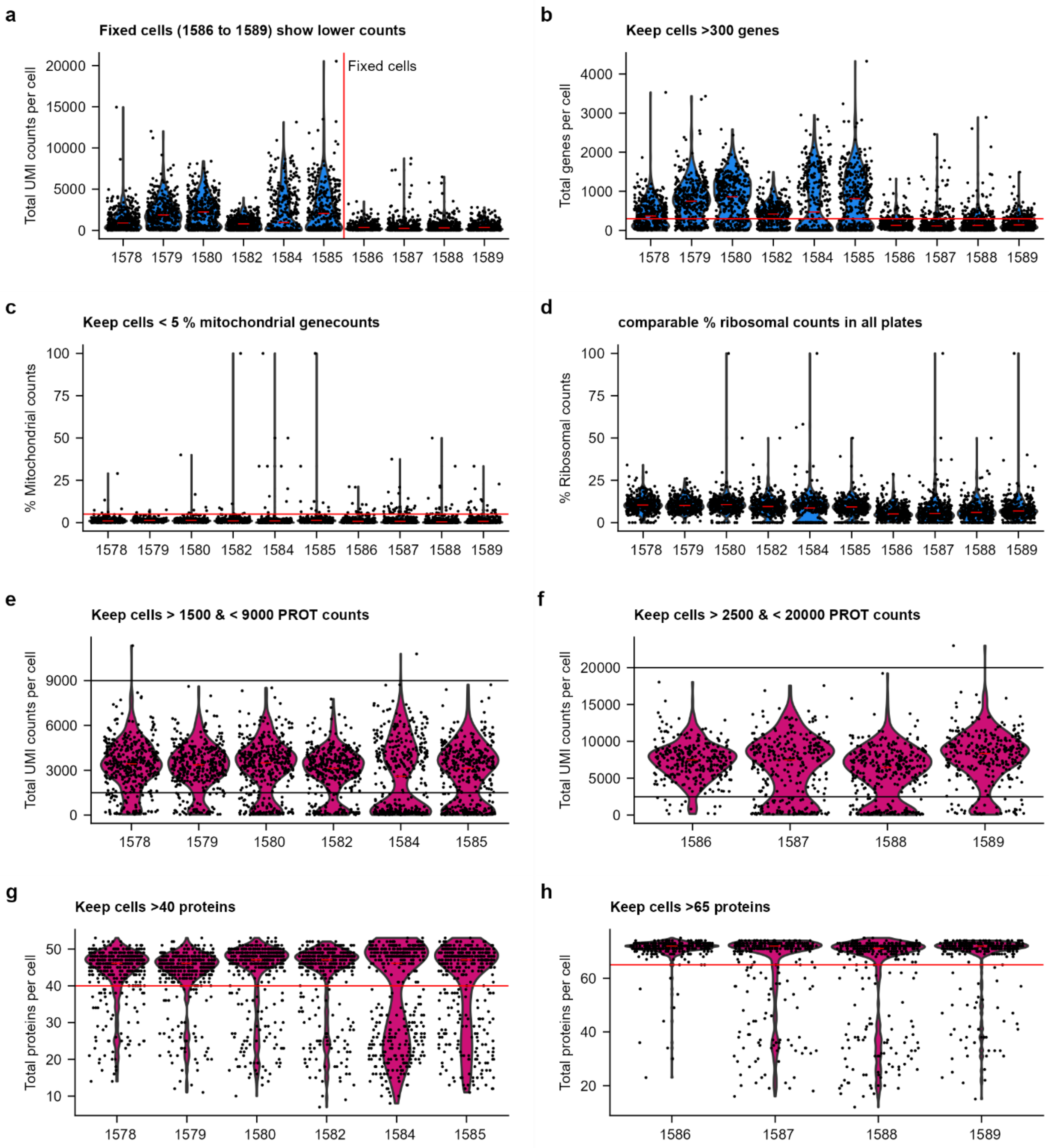


**Fig. S3. Quality control of single-cell RNA (blue, A-D) and antibody (pink, E-H) libraries.** (**A**) The number of unique molecular identifiers filtered mapped (UMIFM) detected per cell. (**B**) The number of different genes detected per cell. Cells with > 300 genes per cell were kept for further analysis. (**C**) Percentage of mitochondrial UMIFM counts out of the total number of UMIFM counts per cell. Cells with < 5% mitochondrial counts per cell were kept for further analysis. (**D**) Percentage of ribosomal UMIFM counts out of the total number of UMIFM counts per cell. (**E**) The number of proteins counts per cell for the live cell sample. Cells with < 1500 and <9000 protein counts per cell were kept for further analysis. (**F**) The number of protein counts per cell for the fixed cell sample. Cells with < 2500 and <20000 protein counts per cell were kept for further analysis. (**G**) The number of different proteins detected per cell for the live cell sample. Cells with > 40 different proteins detected were kept for further analysis. (**H**) The number of different proteins detected per cell for the fixed cell sample. Cells with > 65 different proteins detected were kept for further analysis

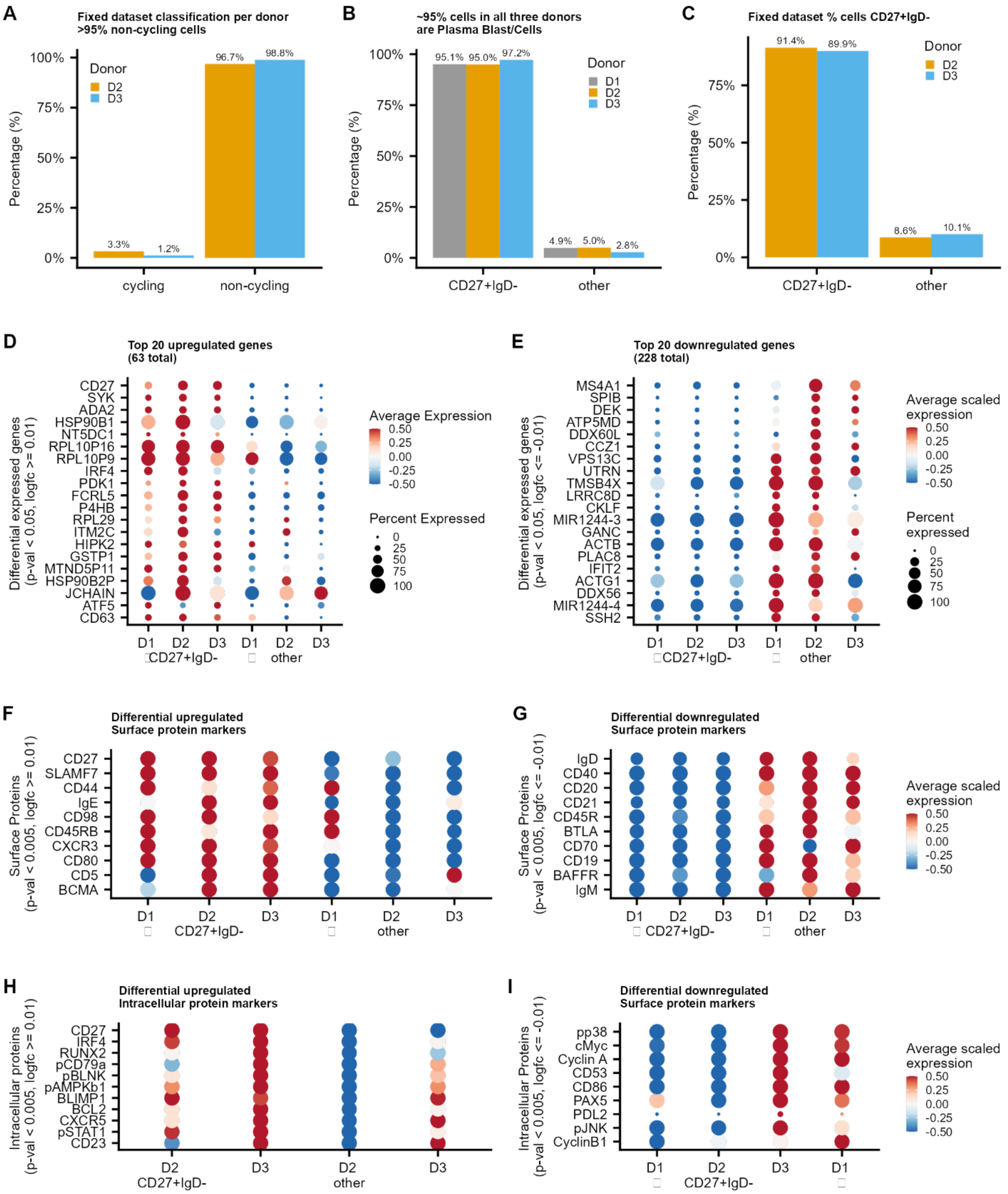


**Fig. S4. Proportions of non-cycling and CD27+IgD- cells and differential genes and proteins.**

(**A**) Percentage of dividing cells in the fixed cell sample per donor. (**B**) Percentage of CD27+IgD- PCs in the live cell sample per donor. (**C**) Percentage of CD27+IgD- PCs in the fixed cell sample per donor. (**D-I**) Differential gene expression of CD27+IgD- gated cells of (**D-E**), Surface proteins (**F-G**) and intracellular (phosphor-)proteins (**H-I**).

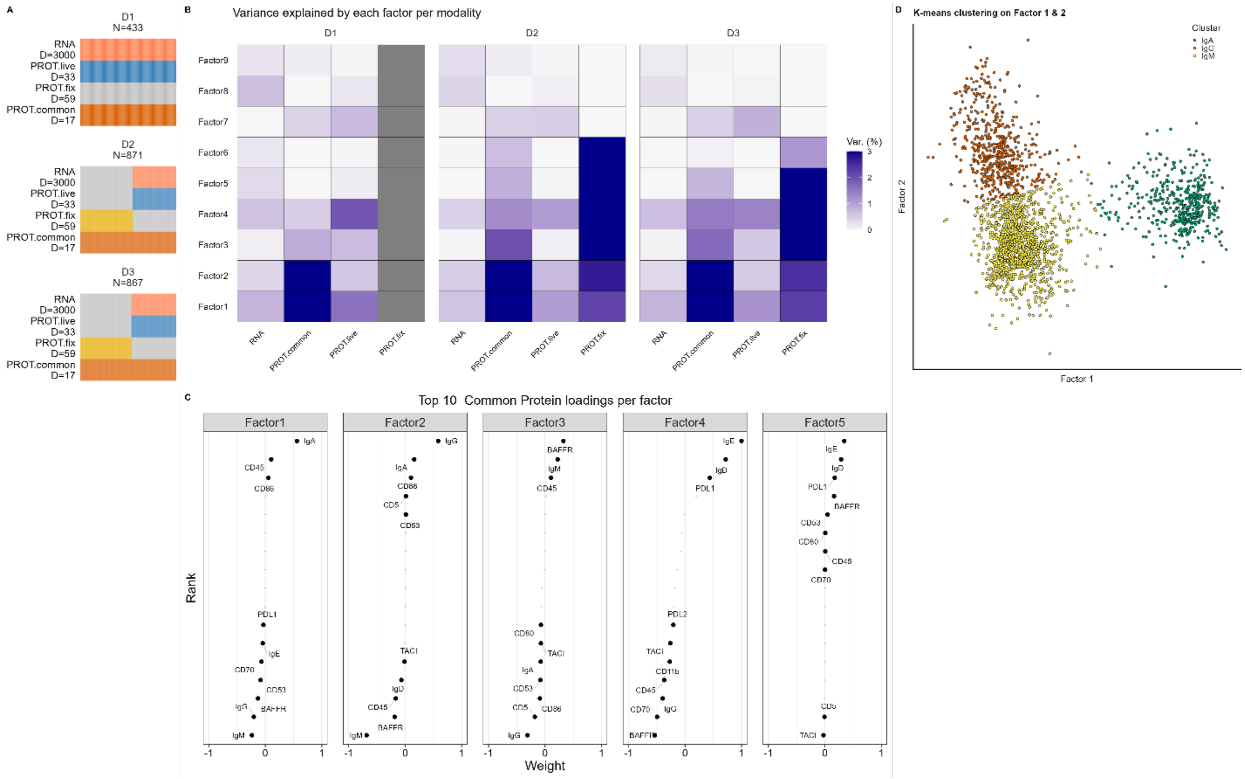


**Fig. S5. Background information of multi-omics factor analysis (MOFA).**

(**A**) Input for MOFA model. Grouped per donor (D1-3), with modalities split into RNA, proteins unique to live-cells, proteins unique to fixed cells and proteins common in both fixed and live cells

(**B**) Percentage variance explained per modality per factor.

(**C**) Top loadings of Factor 1 – 5.

(**D**) K-means clustering (3 clusters) using factor 1 & 2.

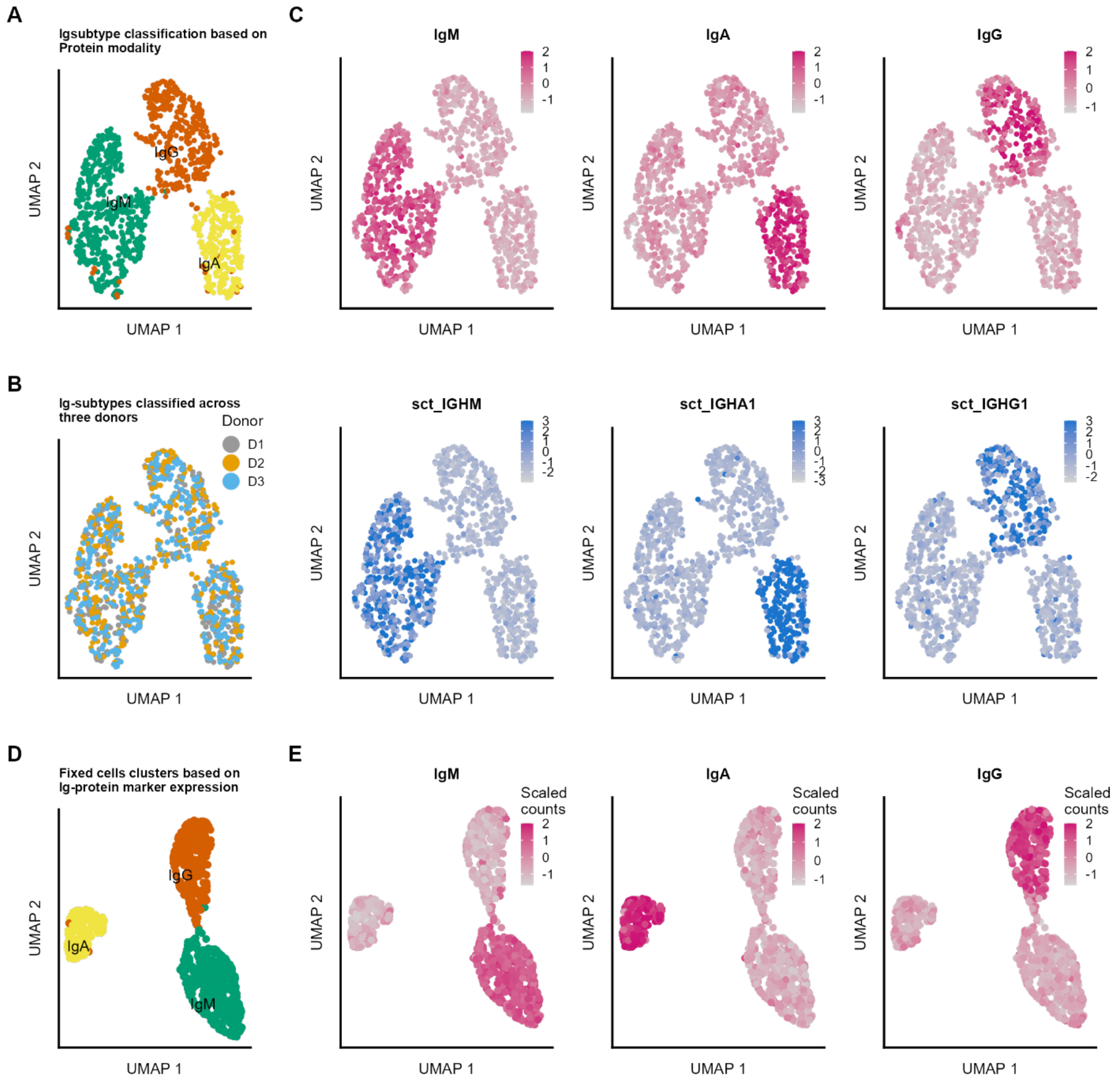


**Fig. S6. Per-dataset analysis of Ig-classes using surface-expression levels.**

(**A**) UMAP representation and clustering based on Ig-protein measurements of the live-cell dataset.
(**B**) UMAP coloured by donor shows all three donors evenly spread over Ig-classes.
(**C**) Protein-levels (upper panels, pink color) and gene-expression (lower panels, blue color) of IgM, IgA and IgG.
(**D**) UMAP representation of PCA performed with Ig-proteins as input of fixed cell dataset. Clustering performed on same principal components.
(**E**) Normalized and scaled protein expression of IgM, IgA or IgG proteins overlaid on UMAP presentation from D.

A

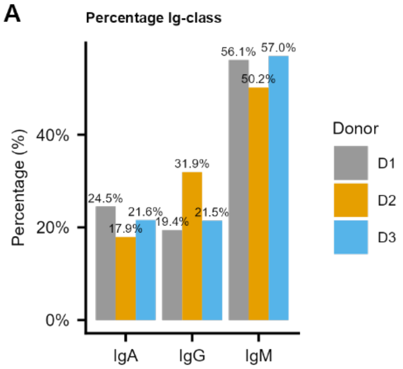


**Fig. S7. Percentage cells IgA, IgM and IgG.** For three donors the percentage of IgA, IgG or IgM classified cells are shown.

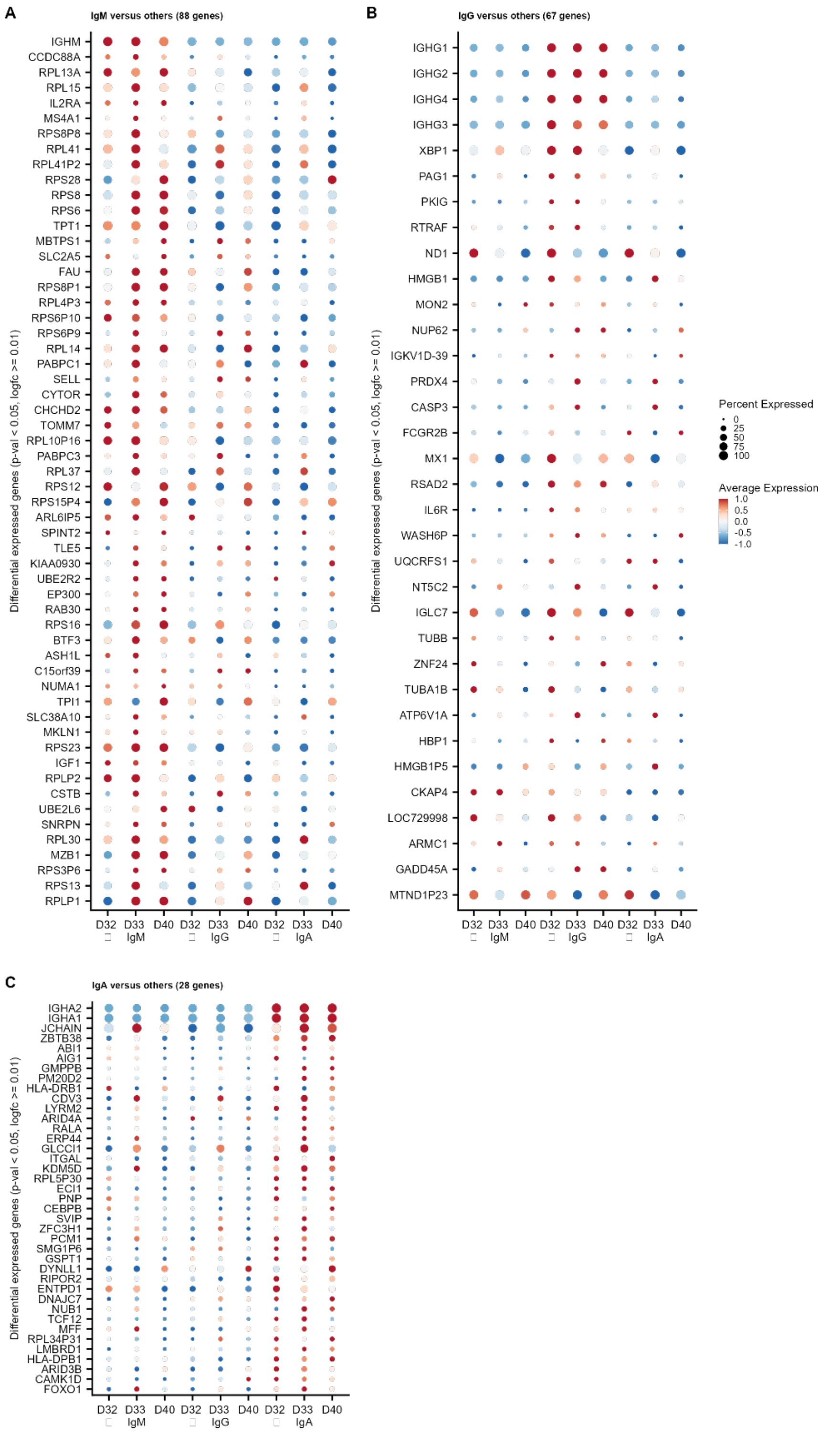


**Fig. S8. Differential expressed genes per Ig-class IgM (A), IgG (B), IgA (C)**

**Table S1.**

Antibody panel with DNA barcodes used on fixed cells. In green highlighted overlapping antibodies with the live-cell panel (Table S2, same clone numbers).

| | Target | Ab barcode sequence | Vendor | Clone | Used concentration |
|---|---|---|---|---|---|
| 1 | **Histone H3** | **CAATCCCT** | Cell Signaling 4499BF | D1H2 | 1 µg/ml |
| 2 | **p-p65** | **GTCCAGGC** | Cell Signaling 3033BF | 93H1 | 1 µg/ml |
| 3 | **p-SYK** | **TGTGTATA** | Cell Signaling 2710BF | C87C1 | 1 µg/ml |
| 4 | **p-JNK** | **AGGATCGA** | Cell Signaling 9255BF | G9 | 1 µg/ml |
| 5 | **p-p38** | **CACGATTC** | Cell Signaling 4511BF | D3F9 | 1 µg/ml |
| 6 | **BCL2** | **GTATCGAG** | Biolegend 658702 | 100 | 1 µg/ml |
| 7 | **TACI** | **TCTCGACT** | Biolegend 311902 | 1A1 | 1 µg/ml |
| 8 | **CD138** | **CATGCGTA** | Abcam ab216458 | EPR6454 | 1 µg/ml |
| 9 | **Cyclin E** | **GTGATAGT** | ThermoFisher 32-1600 | HE12 | 1 µg/ml |
| 10 | **c-MYC** | **TGATATCG** | ThermoFisher 700648 | 27H46L35 | 1 µg/ml |
| 11 | **p-RB** | **AGGGCGTT** | ThermoFisher 701059 | 14H7L14 | 1 µg/ml |
| 12 | **p-c-JUN** | **CTATACGC** | ThermoFisher MA5-27760 | GT653) | 1 µg/ml |
| 13 | **CD70** | **TACATAAG** | BD Bioscience 555833 | Ki-24 | 1 µg/ml |
| 14 | **IgD** | **AATTGAAC** | BD Bioscience 555776 | IA6-2 | 1 µg/ml |
| 15 | **CD86** | **CCAGTGGA** | BD Bioscience 555655 | 2331 (FUN-1) | 1 µg/ml |
| 16 | **CD20** | **GTCCATTG** | BD Bioscience 555677 | H1 | 1 µg/ml |
| 17 | **CD79a** | **TGGACCCT** | BD Bioscience 555934 | HM47 | 1 µg/ml |
| 19 | **CD5** | **AGCAGTTA** | BD Bioscience 555350 | UCHT2 | 1 µg/ml |
| 19 | **PAX5** | **CTTGTACC** | Biolegend 649702 | 1H9 | 1 µg/ml |
| 20 | **PD-L1** | **GAACCCGG** | R&D Systems MAB1561 | 130021 | 1 µg/ml |
| 21 | **PD-L2** | **TCGTAGAT** | R&D Systems MAB1224 | 176611 | 1 µg/ml |
| 22 | **CD21** | **ACGCGGAA** | BD Bioscience 555421 | B-ly4 | 1 µg/ml |
| 23 | **TLR9** | **CGCTATCC** | Biolegend 394802 | S16013D | 1 µg/ml |
| 24 | **TLR10** | **GTTGCATG** | Biolegend 354602 | 3C10C5 | 1 µg/ml |
| 25 | **RUNX2** | **ATCGCCAT** | Abcam ab240329 | EPR14334 | 1 µg/ml |
| 26 | **BTLA** | **CATAAAGG** | Abcam ab254287 | EPR22224-271 | 1 µg/ml |
| 27 | **CXCR3** | **TCACGGTA** | Abcam ab64714 | 49801 | 1 µg/ml |
| 28 | **CD19** | **CACTCAAC** | SantaCruz sc-373897 | F-3 | 1 µg/ml |
| 29 | **p-Histon H3** | **GCTGTGA** | Biolegend 650802 | 11D8 | 1 µg/ml |
| 30 | **IgM** | **TTGCGTCG** | Biolegend 314502 | MHM-88 | 1 µg/ml |
| 31 | **Cyclin A** | **ATATGAGA** | Biolegend 644001 | E23.1 | 1 µg/ml |
| 32 | **p-Histon H2A.X** | **CACCTCAG** | Biolegend 613402 | 2F3 | 1 µg/ml |
| 33 | **Cyclin B1** | **GCTACTTC** | Biolegend 647902 | V152 | 1 µg/ml |
| 34 | **Ki-67** | **TGGGAGCT** | Biolegend 350523 | Ki-67 | 1 µg/ml |
| 35 | **CCR9** | **ATCCGGCA** | Abcam ab247269 | E99 | 1 µg/ml |
| 36 | **p-SRC** | **GGTAATGT** | R&D Systems MAB2685 | 1246F | 1 µg/ml |
| 37 | **p-TOR** | **TAAGCCAC** | R&D Systems MAB1665 | 834115 | 1 µg/ml |

| | | | | | |
|---|---|---|---|---|---|
| 38 | **GAPDH** | **ACCGAACA** | Biolegend 607902 | W17079A | 1 µg/ml |
| 39 | **CD11b** | **GTTTGTGG** | Biolegend 301302 | ICRF44 | 1 µg/ml |
| 40 | **p-BLNK** | **TAGACGAC** | BD Biosciences 558366 | J117-1278 | 1 µg/ml |
| 41 | **CD36** | **ACGCTTGG** | Abcam ab255331 | EPR22509-40 | 1 µg/ml |
| 42 | **CD53** | **CGCTACAT** | BD Biosciences 555506 | HI29 | 1 µg/ml |
| 43 | **CD45** | **TTTGCGTC** | Biolegend 304045 | HI30 | 0.1 µg/ml |
| 44 | **CXCR4** | **ATGGTCGC** | Abcam ab197203 | UMB2 | 1 µg/ml |
| 45 | **p-CD79a** | **CGACATAG** | Cell Signaling 14732BF | D1B9 | 1 µg/ml |
| 46 | **CXCR5** | **GATTCGCT** | Abcam ab225575 | EPR8837 | 1 µg/ml |
| 47 | **CD95** | **TCCAGATA** | Abcam ab227907 | EPR21088 | 1 µg/ml |
| 48 | **Integrin β1** | **ACTACTGT** | Thermo Fisher 700241 | 9H26L42 | 1 µg/ml |
| 49 | **p-AMPK-b1** | **CGGGAACG** | Thermo Fisher 700241 | 9H26L42 | 1 µg/ml |
| 50 | **T-bet** | **GACCTCTC** | Biolegend 644825 | 4B10 | 1 µg/ml |
| 51 | **p-IKK a/b** | **TTATGGAA** | ThermoFisher 701643 | 7H17L17 | 1 µg/ml |
| 52 | **CD27** | **ACAGCAAC** | Abcam ab192336 | EPR8569 | 1 µg/ml |
| 53 | **p-JAK1** | **CGCAATTT** | ThermoFisher 700028 | 59H4L5 | 1 µg/ml |
| 54 | **Integrin β7** | **GAGTTGCG** | Abcam ab246697 | EP5948 | 1 µg/ml |
| 55 | **IL10** | **TTTCGCGA** | ThermoFisher 16-7108-85 | JES3-9D7 | 1 µg/ml |
| 56 | **CCL3/4** | **ACCAGTCC** | R&D Systems MAB2701-100 | 93342 | 1 µg/ml |
| 57 | **CD80** | **CTTTCCTT** | Biolegend 305212 | 2D10 | 1 µg/ml |
| 58 | **Integrin α4** | **CTGGACGT** | Abcam ab22858 | HP2/1 | 1 µg/ml |
| 59 | **p-STAT1** | **GAACGGTC** | ThermoFisher 33-3400 | ST1P-11A5 | 1 µg/ml |
| 60 | **p-STAT3** | **TGTTCACG** | Biolegend 690402 | A16002B | 1 µg/ml |
| 61 | **p-STAT5** | **ATCTGATC** | ThermoFisher 701063 | 6H5L15 | 1 µg/ml |
| 62 | **p-STAT6** | **GAGAAGGG** | ThermoFisher 700247 | 46H1L12 | 1 µg/ml |
| 63 | **KLF6** | **TCACTCCT** | ThermoFisher 39-6900 | 9A2 | 1 µg/ml |
| 64 | **BCL6** | **AGATAACA** | BD Bioscneces 561520 | K112-91 | 1 µg/ml |
| 65 | **AID** | **CTTATTTG** | BD Biosciences 565784 | EK2-5G9 | 1 µg/ml |
| 66 | **IgG** | **GCGGGCAT** | BD Biosciences 555784 | G18-145 | 1 µg/ml |
| 67 | **CD24** | **TACCCGGC** | RnD Systems MAB5247 | ML5 | 1 µg/ml |
| 68 | **IgA** | **ATTGTTTC** | Biolegend 411502 | HP6123 | 1 µg/ml |
| 69 | **IgE** | **CGCAGGAG** | Biolegend 325502 | MHE-18 | 1 µg/ml |
| 70 | **CD23** | **GCACCAGT** | Abcam ab245732 | SP163 | 1 µg/ml |
| 71 | **BLIMP1** | **TAGTACCA** | RnD Systems MAB36081 | 646702 | 1 µg/ml |
| 72 | **IRF8** | **ATTCGTGC** | Biolegend 656502 | 656502 | 1 µg/ml |
| 73 | **IRF4** | **CGCGTGCA** | ThermoFisher 14-9858-82 | 3 E 4 | 1 µg/ml |
| 74 | **BAFFR** | **GAATACTG** | RnD Systems MAB1162 | 2403C | 1 µg/ml |
| 75 | **XBP1** | **TCGACAAT** | Abcam ab239954 | EPR4086 | 1 µg/ml |

**Table S2.**

Antibody panel with DNA barcodes used on non-fixed cells.

| | Target | Ab barcode sequence | Vendor | Clone | Used concentration |
|---|---|---|---|---|---|
| 1 | **CD19** | **CAATCCCT** | Biolegend 302201 | HIB19 | 1 µg/ml |
| 2 | **CD20** | **GTCCAGGC** | Biolegend 302301 | 2H7 | 1 µg/ml |
| 3 | **CD22** | **TGTGTATA** | Biolegend 302502 | HIB22 | 1 µg/ml |
| 4 | **CD25** | **AGGATCGA** | Biolegend 302602 | BC96 | 1 µg/ml |
| 5 | **CD27** | **CACGATTC** | Biolegend 356401 | M-T271 | 1 µg/ml |
| 6 | **TACI** | **TCTCGACT** | Biolegend 311902 | 1A1 | 1 µg/ml |
| 7 | **CD31** | **ACCCGCAC** | Biolegend 303101 | WM59 | 1 µg/ml |
| 8 | **CD32** | **CATGCGTA** | Biolegend 303201 | FUN-2 | 1 µg/ml |
| 9 | **CD36** | **GTGATAGT** | Biolegend 336202 | 5-271 | 1 µg/ml |
| 10 | **CD40** | **TGATATCG** | Biolegend 334302 | 5C3 | 1 µg/ml |
| 11 | **CD44** | **AGGGCGTT** | Biolegend 103001 | IM7 | 1 µg/ml |
| 12 | **CD45RB** | **CTATACGC** | Biolegend 310202 | MEM-55 | 1 µg/ml |
| 13 | **CD49d** | **GCTCGTCA** | Biolegend 304301 | 9F10 | 1 µg/ml |
| 14 | **CD70** | **TACATAAG** | BD Bioscience 555833 | Ki-24 | 1 µg/ml |
| 15 | **IgD** | **AATTGAAC** | BD Bioscience 555776 | IA6-2 | 1 µg/ml |
| 16 | **CD86** | **CCAGTGGA** | BD Bioscience 555655 | 2331 (FUN-1) | 1 µg/ml |
| 17 | **CD79a** | **TGGACCCT** | Biolegend 333502 | HM47 | 1 µg/ml |
| 18 | **CD5** | **AGCAGTTA** | BD Bioscience 555350 | UCHT2 | 1 µg/ml |
| 19 | **C138** | **CTTGTACC** | Biolegend 356501 | MI15 | 1 µg/ml |
| 20 | **C200** | **GAACCCGG** | Biolegend 329201 | OX-104 | 1 µg/ml |
| 21 | **BTLA** | **TCGTAGAT** | Biolegend 344502 | MIH26 | 1 µg/ml |
| 22 | **CD21** | **ACGCGGAA** | BD Bioscience 555421 | B-ly4 | 1 µg/ml |
| 23 | **BCMA** | **CGCTATCC** | Biolegend 357502 | 19F2 | 1 µg/ml |
| 24 | **CCR9** | **GTTGCATG** | Biolegend 358902 | L053E8 | 1 µg/ml |
| 25 | **CCR10** | **TAAATCGT** | Biolegend 341502 | 6588-5 | 1 µg/ml |
| 26 | **CXCR3** | **ATCGCCAT** | Biolegend 353702 | G025H7 | 1 µg/ml |
| 27 | **CXCR5** | **TCACGGTA** | Biolegend 356901 | J252D4 | 1 µg/ml |
| 28 | **CD45R** | **CACTCAAC** | Biolegend 103201 | RA3-6B2 | 1 µg/ml |
| 29 | **IL6R** | **GCTGTGA** | Biolegend 352801 | UV4 | 1 µg/ml |
| 30 | **IgM** | **TTGCGTCG** | Biolegend 314502 | MHM-88 | 1 µg/ml |
| 31 | **Integrin β7** | **ATATGAGA** | Biolegend 321202 | FIB504 | 1 µg/ml |
| 32 | **CD98** | **CACCTCAG** | Biolegend 315602 | MEM-108 | 1 µg/ml |
| 33 | **SLAMF7** | **GCTACTTC** | Biolegend 331802 | 162.1 | 1 µg/ml |
| 34 | **CCR7** | **TGGGAGCT** | Biolegend 353202 | G043H7 | 1 µg/ml |
| 35 | **LAG3** | **ATCCGGCA** | Biolegend 369302 | 11C3C65 | 1 µg/ml |
| 36 | **LT-βR** | **CCGTTATG** | Biolegend 322002 | 31G4D8 | 1 µg/ml |
| 37 | **CD83** | **GGTAATGT** | RnD Systems MAB1774 | HB15e | 1 µg/ml |
| 38 | **Integrin β1** | **TAAGCCAC** | Abcam ab24693 | P5D2 | 1 µg/ml |
| 39 | **CD56** | **ACCGAACA** | Biolegend 304602 | MEM-188 | 1 µg/ml |
| 40 | **CD11b** | **CGACTCTT** | Biolegend 301302 | ICRF44 | 1 µg/ml |

| | | | | | |
|---|---|---|---|---|---|
| 41 | **PD-L1** | **GTTTGTGG** | RnD Systems MAB1561 | 130021 | 1 µg/ml |
| 42 | **PD-L2** | **TAGACGAC** | RnD Systems MAB1224 | 176611 | 1 µg/ml |
| 43 | **CD53** | **CGCTACAT** | BD Biosciences 555506 | HI29 | 1 µg/ml |
| 44 | **CD45** | **TTTGCGTC** | Biolegend 304045 | HI30 | 0.1 µg/ml |
| 45 | **PD1** | **CGACATAG** | RnD Systems MAB10863 | 2335A | 1 µg/ml |
| 46 | **CD80** | **CTTTCCTT** | Biolegend 305212 | 2D10 | 1 µg/ml |
| 47 | **IgG** | **GCGGGCAT** | BD Biosciences 555784 | G18-145 | 1 µg/ml |
| 48 | **CD24** | **TACCCGGC** | RnD Systems MAB5247 | ML5 | 1 µg/ml |
| 49 | **IgA** | **ATTGTTTC** | Biolegend 411502 | HP6123 | 1 µg/ml |
| 50 | **IgE** | **CGCAGGAG** | Biolegend 325502 | MHE-18 | 1 µg/ml |
| 51 | **CD23** | **GCACCAGT** | Abcam ab245732 | SP163 | 1 µg/ml |
| 52 | **BAFF-R** | **GAATACTG** | RnD Systems MAB1162 | 2403C | 1 µg/ml |

**Table S3.**

Oligo sequences coupled to antibodies from fixed panel

| | Full oligo Sequence (Bold Barcode) | Coupled antibody |
|---|---|---|
| 1 | TCGTCGGCAGCGTCAGATGTGTATAAGAGACAG**CAATCCCT**<br>AAAAAAAAAAAAAAAAAAAAAAAAA | **HistoneH3** |
| 2 | TCGTCGGCAGCGTCAGATGTGTATAAGAGACAG**GTCCAGGC**<br>AAAAAAAAAAAAAAAAAAAAAAAAA | **pp65** |
| 3 | TCGTCGGCAGCGTCAGATGTGTATAAGAGACAG**TGTGTATA**<br>AAAAAAAAAAAAAAAAAAAAAAAAA | **pSyk** |
| 4 | TCGTCGGCAGCGTCAGATGTGTATAAGAGACAG**AGGATCG**<br>**A**AAAAAAAAAAAAAAAAAAAAAAAAA | **pJNK** |
| 5 | TCGTCGGCAGCGTCAGATGTGTATAAGAGACAG**CACGATTC**<br>AAAAAAAAAAAAAAAAAAAAAAAAA | **pp38** |
| 6 | TCGTCGGCAGCGTCAGATGTGTATAAGAGACAG**GTATCGAG**<br>AAAAAAAAAAAAAAAAAAAAAAAAA | **BCL2** |
| 7 | TCGTCGGCAGCGTCAGATGTGTATAAGAGACAG**TCTCGACT**<br>AAAAAAAAAAAAAAAAAAAAAAAAA | **TACI** |
| 8 | TCGTCGGCAGCGTCAGATGTGTATAAGAGACAG**CATGCGTA**<br>AAAAAAAAAAAAAAAAAAAAAAAAA | **CD138** |
| 9 | TCGTCGGCAGCGTCAGATGTGTATAAGAGACAG**GTGATAGT**<br>AAAAAAAAAAAAAAAAAAAAAAAAA | **CyclinE** |
| 10 | TCGTCGGCAGCGTCAGATGTGTATAAGAGACAG**TGATATCG**<br>AAAAAAAAAAAAAAAAAAAAAAAAA | **cMyc** |
| 11 | TCGTCGGCAGCGTCAGATGTGTATAAGAGACAG**AGGGCGTT**<br>AAAAAAAAAAAAAAAAAAAAAAAAA | **p-Rb** |
| 12 | TCGTCGGCAGCGTCAGATGTGTATAAGAGACAG**CTATACGC**<br>AAAAAAAAAAAAAAAAAAAAAAAAA | **p-c-Jun** |
| 13 | TCGTCGGCAGCGTCAGATGTGTATAAGAGACAG**TACATAAG**<br>AAAAAAAAAAAAAAAAAAAAAAAAA | **CD70** |
| 14 | TCGTCGGCAGCGTCAGATGTGTATAAGAGACAG**AATTGAAC**<br>AAAAAAAAAAAAAAAAAAAAAAAAA | **IgD** |
| 15 | TCGTCGGCAGCGTCAGATGTGTATAAGAGACAG**CCAGTGGA**<br>AAAAAAAAAAAAAAAAAAAAAAAAA | **CD86** |
| 16 | TCGTCGGCAGCGTCAGATGTGTATAAGAGACAG**GTCCATTG**<br>AAAAAAAAAAAAAAAAAAAAAAAAA | **CD20** |
| 17 | TCGTCGGCAGCGTCAGATGTGTATAAGAGACAG**TGGACCCT**<br>AAAAAAAAAAAAAAAAAAAAAAAAA | **CD79a** |
| 19 | TCGTCGGCAGCGTCAGATGTGTATAAGAGACAG**AGCAGTTA**<br>AAAAAAAAAAAAAAAAAAAAAAAAA | **CD5** |
| 19 | TCGTCGGCAGCGTCAGATGTGTATAAGAGACAG**CTTGTACC**<br>AAAAAAAAAAAAAAAAAAAAAAAAA | **PAX5** |
| 20 | TCGTCGGCAGCGTCAGATGTGTATAAGAGACAG**GAACCCGG**<br>AAAAAAAAAAAAAAAAAAAAAAAAA | **PDL1** |
| 21 | TCGTCGGCAGCGTCAGATGTGTATAAGAGACAG**TCGTAGAT**<br>AAAAAAAAAAAAAAAAAAAAAAAAA | **PDL2** |

| | | |
|---|---|---|
| 22 | TCGTCGGCAGCGTCAGATGTGTATAAGAGACAG**ACGCGGA**<br>**A**AAAAAAAAAAAAAAAAAAAAAAAAA | **CD21** |
| 23 | TCGTCGGCAGCGTCAGATGTGTATAAGAGACAG**CGCTATCC**<br>AAAAAAAAAAAAAAAAAAAAAAAAA | **TLR9** |
| 24 | TCGTCGGCAGCGTCAGATGTGTATAAGAGACAG**GTTGCATG**<br>AAAAAAAAAAAAAAAAAAAAAAAAA | **TLR10** |
| 25 | TCGTCGGCAGCGTCAGATGTGTATAAGAGACAG**ATCGCCAT**<br>AAAAAAAAAAAAAAAAAAAAAAAAA | **RUNX2** |
| 26 | TCGTCGGCAGCGTCAGATGTGTATAAGAGACAG**CATAAAGG**<br>AAAAAAAAAAAAAAAAAAAAAAAAA | **BTLA** |
| 27 | TCGTCGGCAGCGTCAGATGTGTATAAGAGACAG**TCACGGTA**<br>AAAAAAAAAAAAAAAAAAAAAAAAA | **CXCR3** |
| 28 | TCGTCGGCAGCGTCAGATGTGTATAAGAGACAG**CACTCAAC**<br>AAAAAAAAAAAAAAAAAAAAAAAAA | **CD19** |
| 29 | TCGTCGGCAGCGTCAGATGTGTATAAGAGACAG**GCTGTGA**A<br>AAAAAAAAAAAAAAAAAAAAAAAA | **pHiston H3** |
| 30 | TCGTCGGCAGCGTCAGATGTGTATAAGAGACAG**TTGCGTCG**<br>AAAAAAAAAAAAAAAAAAAAAAAAA | **IgM** |
| 31 | TCGTCGGCAGCGTCAGATGTGTATAAGAGACAG**ATATGAGA**<br>AAAAAAAAAAAAAAAAAAAAAAAAA | **Cyclin A** |
| 32 | TCGTCGGCAGCGTCAGATGTGTATAAGAGACAG**CACCTCAG**<br>AAAAAAAAAAAAAAAAAAAAAAAAA | **pHistonH2AX** |
| 33 | TCGTCGGCAGCGTCAGATGTGTATAAGAGACAG**GCTACTTC**<br>AAAAAAAAAAAAAAAAAAAAAAAAA | **CyclinB1** |
| 34 | TCGTCGGCAGCGTCAGATGTGTATAAGAGACAG**TGGGAGCT**<br>AAAAAAAAAAAAAAAAAAAAAAAAA | **Ki67** |
| 35 | TCGTCGGCAGCGTCAGATGTGTATAAGAGACAG**ATCCGGCA**<br>AAAAAAAAAAAAAAAAAAAAAAAAA | **CCR9** |
| 36 | TCGTCGGCAGCGTCAGATGTGTATAAGAGACAG**GGTAATGT**<br>AAAAAAAAAAAAAAAAAAAAAAAAA | **pSrc** |
| 37 | TCGTCGGCAGCGTCAGATGTGTATAAGAGACAG**TAAGCCAC**<br>AAAAAAAAAAAAAAAAAAAAAAAAA | **pTOR** |
| 38 | TCGTCGGCAGCGTCAGATGTGTATAAGAGACAG**ACCGAACA**<br>AAAAAAAAAAAAAAAAAAAAAAAAA | **GAPDH** |
| 39 | TCGTCGGCAGCGTCAGATGTGTATAAGAGACAG**CGACTCTT**<br>AAAAAAAAAAAAAAAAAAAAAAAAA | **CD11b** |
| 40 | TCGTCGGCAGCGTCAGATGTGTATAAGAGACAG**TAGACGAC**<br>AAAAAAAAAAAAAAAAAAAAAAAAA | **pBLNK** |
| 41 | TCGTCGGCAGCGTCAGATGTGTATAAGAGACAG**ACGCTTGG**<br>AAAAAAAAAAAAAAAAAAAAAAAAA | **CD36** |
| 42 | TCGTCGGCAGCGTCAGATGTGTATAAGAGACAG**CGCTACAT**<br>AAAAAAAAAAAAAAAAAAAAAAAAA | **CD53** |
| 43 | TCGTCGGCAGCGTCAGATGTGTATAAGAGACAG**GAAAGAC**<br>**A**AAAAAAAAAAAAAAAAAAAAAAAAA | **CD38** |
| 44 | TCGTCGGCAGCGTCAGATGTGTATAAGAGACAG**TTTGCGTC**<br>AAAAAAAAAAAAAAAAAAAAAAAAA | **CD45** |

| | | |
|---|---|---|
| 45 | TCGTCGGCAGCGTCAGATGTGTATAAGAGACAG**ATGGTCGC**AAAAAAAAAAAAAAAAAAAAAAAAA | **CXCR4** |
| 46 | TCGTCGGCAGCGTCAGATGTGTATAAGAGACAG**CGACATAG**AAAAAAAAAAAAAAAAAAAAAAAAA | **pCD79a** |
| 47 | TCGTCGGCAGCGTCAGATGTGTATAAGAGACAG**GATTCGCT**AAAAAAAAAAAAAAAAAAAAAAAAA | **CXCR5** |
| 48 | TCGTCGGCAGCGTCAGATGTGTATAAGAGACAG**TCCAGATA**AAAAAAAAAAAAAAAAAAAAAAAAA | **CD95** |
| 49 | TCGTCGGCAGCGTCAGATGTGTATAAGAGACAG**ACTACTGT**AAAAAAAAAAAAAAAAAAAAAAAAA | **IntegrinB1** |
| 50 | TCGTCGGCAGCGTCAGATGTGTATAAGAGACAG**CGGGAACG**AAAAAAAAAAAAAAAAAAAAAAAAA | **pAMPKb1** |
| 51 | TCGTCGGCAGCGTCAGATGTGTATAAGAGACAG**GACCTCTC**AAAAAAAAAAAAAAAAAAAAAAAAA | **Tbet** |
| 52 | TCGTCGGCAGCGTCAGATGTGTATAAGAGACAG**TTATGGAA**AAAAAAAAAAAAAAAAAAAAAAAAA | **pIKKab** |
| 53 | TCGTCGGCAGCGTCAGATGTGTATAAGAGACAG**ACAGCAAC**AAAAAAAAAAAAAAAAAAAAAAAAA | **CD27** |
| 54 | TCGTCGGCAGCGTCAGATGTGTATAAGAGACAG**CGCAATTT**AAAAAAAAAAAAAAAAAAAAAAAAA | **pJAK1** |
| 55 | TCGTCGGCAGCGTCAGATGTGTATAAGAGACAG**GAGTTGCG**AAAAAAAAAAAAAAAAAAAAAAAAA | **IntegrinB7** |
| 56 | TCGTCGGCAGCGTCAGATGTGTATAAGAGACAG**TTTCGCGA**AAAAAAAAAAAAAAAAAAAAAAAAA | **IL10** |
| 57 | TCGTCGGCAGCGTCAGATGTGTATAAGAGACAG**ACCAGTCC**AAAAAAAAAAAAAAAAAAAAAAAAA | **CCL3** |
| 58 | TCGTCGGCAGCGTCAGATGTGTATAAGAGACAG**CTTTCCTT**AAAAAAAAAAAAAAAAAAAAAAAAA | **CD80** |
| 59 | TCGTCGGCAGCGTCAGATGTGTATAAGAGACAG**CTGGACGT**AAAAAAAAAAAAAAAAAAAAAAAAA | **IntegrinA4** |
| 60 | TCGTCGGCAGCGTCAGATGTGTATAAGAGACAG**GAACGGTC**AAAAAAAAAAAAAAAAAAAAAAAAA | **pSTAT1** |
| 61 | TCGTCGGCAGCGTCAGATGTGTATAAGAGACAG**TGTTCACG**AAAAAAAAAAAAAAAAAAAAAAAAA | **pSTAT3** |
| 62 | TCGTCGGCAGCGTCAGATGTGTATAAGAGACAG**ATCTGATC**AAAAAAAAAAAAAAAAAAAAAAAAA | **pSTAT5** |
| 63 | TCGTCGGCAGCGTCAGATGTGTATAAGAGACAG**GAGAAGGG**AAAAAAAAAAAAAAAAAAAAAAAAA | **pSTAT6** |
| 64 | TCGTCGGCAGCGTCAGATGTGTATAAGAGACAG**TCACTCCT**AAAAAAAAAAAAAAAAAAAAAAAAA | **KLF6** |
| 65 | TCGTCGGCAGCGTCAGATGTGTATAAGAGACAG**AGATAACA**AAAAAAAAAAAAAAAAAAAAAAAAA | **Bcl6** |
| 66 | TCGTCGGCAGCGTCAGATGTGTATAAGAGACAG**CTTATTTG**AAAAAAAAAAAAAAAAAAAAAAAAA | **AID** |
| 67 | TCGTCGGCAGCGTCAGATGTGTATAAGAGACAG**GCGGGCAT**AAAAAAAAAAAAAAAAAAAAAAAAA | **IgG** |

| | | |
|---|---|---|
| 68 | TCGTCGGCAGCGTCAGATGTGTATAAGAGACAG**TACCCGGC** AAAAAAAAAAAAAAAAAAAAAAAAA | **CD24** |
| 69 | TCGTCGGCAGCGTCAGATGTGTATAAGAGACAG**ATTGTTTC** AAAAAAAAAAAAAAAAAAAAAAAAA | **IgA** |
| 70 | TCGTCGGCAGCGTCAGATGTGTATAAGAGACAG**CGCAGGA** **G**AAAAAAAAAAAAAAAAAAAAAAAAA | **IgE** |
| 71 | TCGTCGGCAGCGTCAGATGTGTATAAGAGACAG**GCACCAGT** AAAAAAAAAAAAAAAAAAAAAAAAA | **CD23** |
| 72 | TCGTCGGCAGCGTCAGATGTGTATAAGAGACAG**TAGTACCA** AAAAAAAAAAAAAAAAAAAAAAAAA | **BLIMP1** |
| 73 | TCGTCGGCAGCGTCAGATGTGTATAAGAGACAG**ATTCGTGC** AAAAAAAAAAAAAAAAAAAAAAAAA | **IRF8** |
| 74 | TCGTCGGCAGCGTCAGATGTGTATAAGAGACAG**CGCGTGCA** AAAAAAAAAAAAAAAAAAAAAAAAA | **IRF4** |
| 75 | TCGTCGGCAGCGTCAGATGTGTATAAGAGACAG**GAATACTG** AAAAAAAAAAAAAAAAAAAAAAAAA | **BAFFR** |

**Table S4.**

Oligo sequences coupled to antibodies from live-cell panel

| | Full oligo Sequence (Bold Barcode) | Coupled antibody |
|---|---|---|
| 1 | TCGTCGGCAGCGTCAGATGTGTATAAGAGACAG**CAATCCCT**<br>AAAAAAAAAAAAAAAAAAAAAAAAA | **CD19** |
| 2 | TCGTCGGCAGCGTCAGATGTGTATAAGAGACAG**GTCCAGGC**<br>AAAAAAAAAAAAAAAAAAAAAAAAA | **CD20** |
| 3 | TCGTCGGCAGCGTCAGATGTGTATAAGAGACAG**TGTGTATA**<br>AAAAAAAAAAAAAAAAAAAAAAAAA | **CD22** |
| 4 | TCGTCGGCAGCGTCAGATGTGTATAAGAGACAG**AGGATCG**<br>**A**AAAAAAAAAAAAAAAAAAAAAAAAA | **CD25** |
| 5 | TCGTCGGCAGCGTCAGATGTGTATAAGAGACAG**CACGATTC**<br>AAAAAAAAAAAAAAAAAAAAAAAAA | **CD27** |
| 6 | TCGTCGGCAGCGTCAGATGTGTATAAGAGACAG**TCTCGACT**<br>AAAAAAAAAAAAAAAAAAAAAAAAA | **TACI** |
| 7 | TCGTCGGCAGCGTCAGATGTGTATAAGAGACAG**ACCCGCAC**<br>AAAAAAAAAAAAAAAAAAAAAAAAA | **CD31** |
| 8 | TCGTCGGCAGCGTCAGATGTGTATAAGAGACAG**CATGCGTA**<br>AAAAAAAAAAAAAAAAAAAAAAAAA | **CD32** |
| 9 | TCGTCGGCAGCGTCAGATGTGTATAAGAGACAG**GTGATAGT**<br>AAAAAAAAAAAAAAAAAAAAAAAAA | **CD36** |
| 10 | TCGTCGGCAGCGTCAGATGTGTATAAGAGACAG**TGATATCG**<br>AAAAAAAAAAAAAAAAAAAAAAAAA | **CD40** |
| 11 | TCGTCGGCAGCGTCAGATGTGTATAAGAGACAG**AGGGCGTT**<br>AAAAAAAAAAAAAAAAAAAAAAAAA | **CD44** |
| 12 | TCGTCGGCAGCGTCAGATGTGTATAAGAGACAG**CTATACGC**<br>AAAAAAAAAAAAAAAAAAAAAAAAA | **CD45RB** |
| 13 | TCGTCGGCAGCGTCAGATGTGTATAAGAGACAG**GCTCGTCA**<br>AAAAAAAAAAAAAAAAAAAAAAAAA | **CD49d** |
| 14 | TCGTCGGCAGCGTCAGATGTGTATAAGAGACAG**TACATAAG**<br>AAAAAAAAAAAAAAAAAAAAAAAAA | **CD70** |
| 15 | TCGTCGGCAGCGTCAGATGTGTATAAGAGACAG**AATTGAAC**<br>AAAAAAAAAAAAAAAAAAAAAAAAA | **IgD** |
| 16 | TCGTCGGCAGCGTCAGATGTGTATAAGAGACAG**CCAGTGGA**<br>AAAAAAAAAAAAAAAAAAAAAAAAA | **CD86** |
| 17 | TCGTCGGCAGCGTCAGATGTGTATAAGAGACAG**TGGACCCT**<br>AAAAAAAAAAAAAAAAAAAAAAAAA | **CD79a** |
| 19 | TCGTCGGCAGCGTCAGATGTGTATAAGAGACAG**AGCAGTTA**<br>AAAAAAAAAAAAAAAAAAAAAAAAA | **CD5** |
| 19 | TCGTCGGCAGCGTCAGATGTGTATAAGAGACAG**CTTGTACC**<br>AAAAAAAAAAAAAAAAAAAAAAAAA | **CD138** |
| 20 | TCGTCGGCAGCGTCAGATGTGTATAAGAGACAG**GAACCCGG**<br>AAAAAAAAAAAAAAAAAAAAAAAAA | **CD200** |
| 21 | TCGTCGGCAGCGTCAGATGTGTATAAGAGACAG**TCGTAGAT**<br>AAAAAAAAAAAAAAAAAAAAAAAAA | **BTLA** |

| | | |
|---|---|---|
| 22 | TCGTCGGCAGCGTCAGATGTGTATAAGAGACAG**ACGCGGA**<br>**A**AAAAAAAAAAAAAAAAAAAAAAAA | **CD21** |
| 23 | TCGTCGGCAGCGTCAGATGTGTATAAGAGACAG**CGCTATCC**<br>AAAAAAAAAAAAAAAAAAAAAAAA | **BCMA** |
| 24 | TCGTCGGCAGCGTCAGATGTGTATAAGAGACAG**GTTGCATG**<br>AAAAAAAAAAAAAAAAAAAAAAAA | **CCR9** |
| 25 | TCGTCGGCAGCGTCAGATGTGTATAAGAGACAG**TAAATCGT**<br>AAAAAAAAAAAAAAAAAAAAAAAA | **CCR10** |
| 26 | TCGTCGGCAGCGTCAGATGTGTATAAGAGACAG**ATCGCCAT**<br>AAAAAAAAAAAAAAAAAAAAAAAA | **CXCR3** |
| 27 | TCGTCGGCAGCGTCAGATGTGTATAAGAGACAG**TCACGGTA**<br>AAAAAAAAAAAAAAAAAAAAAAAA | **CXCR5** |
| 28 | TCGTCGGCAGCGTCAGATGTGTATAAGAGACAG**CACTCAAC**<br>AAAAAAAAAAAAAAAAAAAAAAAA | **CD45R** |
| 29 | TCGTCGGCAGCGTCAGATGTGTATAAGAGACAG**GCTGTGA**A<br>AAAAAAAAAAAAAAAAAAAAAAA | **IL6R** |
| 30 | TCGTCGGCAGCGTCAGATGTGTATAAGAGACAG**TTGCGTCG**<br>AAAAAAAAAAAAAAAAAAAAAAAA | **IgM** |
| 31 | TCGTCGGCAGCGTCAGATGTGTATAAGAGACAG**ATATGAGA**<br>AAAAAAAAAAAAAAAAAAAAAAAA | **IntegrinB7** |
| 32 | TCGTCGGCAGCGTCAGATGTGTATAAGAGACAG**CACCTCAG**<br>AAAAAAAAAAAAAAAAAAAAAAAA | **CD98** |
| 33 | TCGTCGGCAGCGTCAGATGTGTATAAGAGACAG**GCTACTTC**<br>AAAAAAAAAAAAAAAAAAAAAAAA | **SLAMF7** |
| 34 | TCGTCGGCAGCGTCAGATGTGTATAAGAGACAG**TGGGAGCT**<br>AAAAAAAAAAAAAAAAAAAAAAAA | **CCR7** |
| 35 | TCGTCGGCAGCGTCAGATGTGTATAAGAGACAG**ATCCGGCA**<br>AAAAAAAAAAAAAAAAAAAAAAAA | **LAG3** |
| 36 | TCGTCGGCAGCGTCAGATGTGTATAAGAGACAG**CCGTTATG**<br>AAAAAAAAAAAAAAAAAAAAAAAA | **LT-βR** |
| 37 | TCGTCGGCAGCGTCAGATGTGTATAAGAGACAG**GGTAATGT**<br>AAAAAAAAAAAAAAAAAAAAAAAA | **CD83** |
| 38 | TCGTCGGCAGCGTCAGATGTGTATAAGAGACAG**TAAGCCAC**<br>AAAAAAAAAAAAAAAAAAAAAAAA | **IntegrinB1** |
| 39 | TCGTCGGCAGCGTCAGATGTGTATAAGAGACAG**ACCGAACA**<br>AAAAAAAAAAAAAAAAAAAAAAAA | **CD56** |
| 40 | TCGTCGGCAGCGTCAGATGTGTATAAGAGACAG**CGACTCTT**<br>AAAAAAAAAAAAAAAAAAAAAAAA | **CD11b** |
| 41 | TCGTCGGCAGCGTCAGATGTGTATAAGAGACAG**GTTTGTGG**<br>AAAAAAAAAAAAAAAAAAAAAAAA | **PDL1** |
| 42 | TCGTCGGCAGCGTCAGATGTGTATAAGAGACAG**TAGACGAC**<br>AAAAAAAAAAAAAAAAAAAAAAAA | **PDL2** |

**Table S5.**

Antibodies used for flow cytometry analysis.

| Target | Fluorophore | Vendor | Clone | Used concentration |
|---|---|---|---|---|
| **p-SYK** | PE | Cell Signaling 6485 | C87C1 | 1/50 |
| **p-PLC-y2** | Alexa 647 | BD Bioscience 558498 | K86-689.37 | 1/100 |
| **p-BTK** | PE | Biolegend 601704 | A16128B | 1/50 |
| **p-p38** | Alexa 488 | Cell Signaling 41768S | 3D7 | 1/50 |
| **p-AKT** | PE | Cell Signaling 5315 | D9E | 1/50 |
| **p-ERK 1/2** | PE | Cell Signaling 75765S | D1H6G | 1/50 |
| **p-JNK** | Alexa 647 | Cell Signaling 9257S | G9 | 1/50 |
| **p-p65** | Alexa 647 | Cell Signaling 5733 | 93H1 | 1/50 |
| **p-S6** | PE | Cell Signaling 5316 | D57.2.2E | 1/800 |
| **CD19** | PerCP/Cy5.5 | Biolegend 302230 | HIB19 | 1/100 |
| **IgD** | BV510 | Biolegend 348220 | IA6-2 | 1/50 |
| **CD27** | FITC | BD Bioscience 555440 | M-T271 | 1/25 |
| **aCD20** | APC-H7 | BD Bioscience 560853 | 2H7 | 1/25 |
| **CD38** | PE-Cy7 | eBioscience 250-388-41 | HB7 | 1/2000 |
| **IgA** | PE | Miltenyi 130-113-472 | IS11-8E10 | 1/200 |

### Table S6.

Sequences of all the primers used for library preparation.

| | Primer sequence 5’ -> 3’ |
|---|---|
| **CEL-seq 2 barcoding primers (as in Gerlach et.al. 2019)** | GCCGGTAATACGACTCACTATAGGGGTTCAGACGTGTGCTCTTCCGATCTNNNNNNNNCGTCTAATTTTTTTTTTTTTTTTTTTTTTTTTV |
| **FWR Random Octamer** | CACGACGCTCTTCCGATCTNNNNNNNN |
| **REV unique indexing primer** | CAAGCAGAAGACGGCATACGAGATATCAGTGTGACTGGAGTTCAGACGTGTGCTCTTCCGATC |
| **FWD LP primer** | AATGATACGGCGACCACCGAGATCTACACTCTTTCCCTACACGACGCTCTTCCGATCT |
| **Nextera primer** | AATGATACGGCGACCACCGAGATCTACACGGTCTCGTCGGCAGCGTCAGATG |
